# Human Lung-resident Mucosal-Associated Invariant T cells are Abundant, Express Antimicrobial Proteins, and are Cytokine Responsive

**DOI:** 10.1101/2022.04.28.489781

**Authors:** Erin W. Meermeier, Christina L. Zheng, Jessica G Tran, Shogo Soma, Aneta H. Worley, David I. Weiss, Robert L. Modlin, Gwendolyn Swarbrick, Elham Karamooz, Sharon Khuzwayo, Emily B. Wong, Marielle C. Gold, David M. Lewinsohn

**Affiliations:** Department of Pulmonary and Critical Care Medicine, Oregon Health & Science University, Portland, OR 97239, USA; Department of Medical Informatics and Clinical Epidemiology, Oregon Health & Science University, Portland, OR 97239, USA; VA Portland Health Care System, Portland, OR 97239, USA; David Geffen School of Medicine at UCLA, Los Angeles, CA 90095, USA; Division of Dermatology, Department of Medicine, David Geffen School of Medicine at University of California, Los Angeles, Los Angeles, CA 90095, USA; Africa Health Research Institute, Durban, South Africa; School of Laboratory Medicine and Medical Sciences, University of KwaZulu-Natal, Durban, South Africa; Division of Infectious Diseases, Massachusetts General Hospital, Boston, MA, USA; Harvard Medical School, Boston, MA, USA; Division of Infection and Immunity, University College London, London, UK

**Author notes:** Correspondence: David M. Lewinsohn, Pulmonary and Critical Care Medicine, OHSU, 3181 SW Sam Jackson Park Rd, Mail Code VA R&D 11, Portland, OR 97239, USA.

## Abstract

Mucosal-associated Invariant T (MAIT) cells are an innate-like T cell subset that recognize a broad array of microbial pathogens, including respiratory pathogens. Here we investigate the transcriptional profile of MAIT cells localized to the human lung, and postulate that MAIT cells may play a role in maintaining homeostasis at this mucosal barrier. Using the MR1/5-OP-RU tetramer, we identified MAIT cells and non-MAIT CD8^+^ T cells in lung tissue not suitable for transplant from human donors. We used RNA-sequencing of MAIT cells compared to non-MAIT CD8^+^ T cells to define the transcriptome of MAIT cells in the human lung. We show that, as a population, lung MAIT cells are polycytotoxic, secrete the directly antimicrobial molecule IL-26, express genes associated with persistence, and selectively express cytokine and chemokine-related molecules distinct from other lung-resident CD8^+^ T cells, such as interferon-*γ−* and IL-12-receptors. These data highlight MAIT cells’ predisposition to rapid pro-inflammatory cytokine responsiveness and antimicrobial mechanisms in human lung tissue, concordant with findings of blood-derived counterparts, and support a function for MAIT cells as early sensors in the defense of respiratory barrier function.

## Introduction

The mucosal surface of the lung constitutes one of the most significant portals of entry for pathogenic microbes. Due to the vital nature of its gas exchange function, cellular immune responses in the respiratory tract need to be poised, efficient, and regulated for appropriate defensive functions. Thus, the organization of a regulatory network of immune cells in the respiratory tract plays an important role in maintaining tissue integrity and defending against microbial invasions. Human mucosal-associated invariant T (MAIT) cells are a unique T cell population found in all humans that are characterized by the use of a semi-invariant T cell receptor (TCR) *α*-chain, dependence on the non-classical MHC Class I-related molecule MR1, and their rapid effector function. MAIT cells are a subset of T cells restricted by MR1, and are able to recognize infection by a broad array of pathogens, including respiratory pathogens.

Collectively, *in vivo* models of lung infection link MAIT cells to the early stages of bacterial containment in the respiratory tract and in shaping the ensuing adaptive immune response ^1–5^. MAIT cell accumulation and expansion at the site of infection depended upon local antigen in conjunction with costimulation from bacteria or TLR signals Postnatally, MAIT cells expand rapidly in mice in a manner that depends on translocation of MAIT cell antigens such as 5-OP-RU ^6, 7^. In humans, while MAIT cells are present as effectors in both thymus and cord blood ^8, 9^ the expansion of canonical MAIT cells occurs rapidly, and would be consistent with the hypothesis that expansion of MAIT cells may be associated with the neonatal development of the microbiome. Supporting a role for MAIT cells in human host-defense, an individual, found to have a mutation in *MR1* resulting in the absence of MAIT cells, presented with reoccurring infectious diseases at mucosal surfaces^10^. At present an in-depth analysis into biological pathways and effector functions used by MAIT cells in the human lung has not been performed. Thus, we sought to stringently define functional characteristics of MAIT cells residing in lung tissue.

A striking observation of MAIT cells is their presence at mucosal tissue sites but whether they form a distinctly functional tissue T cell subset remains unclear. MAIT cells were first discovered by their abundance in the gut lamina propria of humans and mice^11^ and are developmentally predisposed to reside in mucosal sites early in life. In support of this, MAIT cells in the thymus, cord blood and peripheral blood express significantly less CD62L, a selectin associated with homing to lymphoid tissues^8^. Moreover, MAIT cells in human 2^nd^ trimester fetuses already express CD45RO and are functionally mature in mucosal tissues, suggesting the development of an early arm of antibacterial mucosal immunity ^12^. A study by Salou et. al, used transcriptomics, parabiosis, and thymocyte adoptive transfer in mice to show that MAIT and NKT cell subsets share a common differentiation program which is associated with tissue residence^13^. Here, in contrast to other organs, the authors noted that MAIT cells in the lung had slower rates of parabiotic exchange. The authors suggest that a preset transcriptional program predisposes MAIT cells to reside in tissues. We found that MAIT cells with mycobacterial reactivity are present in the airways of healthy individuals, and further enriched in bronchoalveolar fluid isolated from patients with active pulmonary tuberculosis^14^. Furthermore, bronchoalveolar MAIT cells from healthy individuals with latent tuberculosis produce less pro-inflammatory cytokines, display higher levels of inhibitory markers, and can express distinct tissue repair genes compared to peripheral MAIT cells^15^. Finally, murine models of bacterial pneumonia have uncovered a key role for MAIT cells in the lungs that proliferate *in situ* as opposed to being recruited from lymph nodes ^16^. Examination of the function of human lung-derived MAIT cells has been underexplored, and we do not yet know whether they form a functionally distinct T cell subset compared to other CD8^+^ lung-resident T cells.

MAIT cells derived from different tissues have a phenotype and function consistent with their anatomic, microbial, and inflammatory microenvironment. Upon stimulation, MAIT cells produce a variety of cytokines that have the potential to promote an antibacterial or anti-fungal response. It has been widely observed that they produce the pro-inflammatory cytokines IFN-*γ* and TNF but also IL-17 and IL-22 in certain tissues and disease states ^12, 17–21^. Analysis of human intestinal MAIT cells shows transcripts but not corresponding proteins of proinflammatory cytokines suggesting a degree of post-transcriptional regulation^22^. MAIT cells also express cytotoxic molecules and are capable of lysing target cells ranging from macrophages to epithelial cells ^23–25^. The cytotoxic potential of MAIT cells can be licensed by bacterial infection, TCR stimulation, or cytokines including IL-7, IL-12, IL-15, and IL-18 ^23, 25–27^. However, the inflammatory state and cytotoxic potential of MAIT cells in the human lung remains unclear.

Here, we sought to address the hypothesis that MAIT cells in the lung have unique transcriptionally-defined effector functions for their critical role in early recognition and control of respiratory pathogens. We took a comprehensive approach to define the transcriptome of human lung-derived MAIT cells compared to paired non-MAIT CD8^+^ T cells. Our analysis reveals MAIT cell’s comprehensive repertoire of cytokine and chemokine receptors including the IL-12 receptor and IFN-*γ* receptor, and expression and induction of cytolytic molecules for their antimicrobial capacity, such as IL-26 and granzyme B. Our findings align with the paradigm of MAIT cells as a lung-enriched cell type armed for specialized defense against initial pathogen encounters.

## Results

### MAIT cells are enriched in the lungs

To identify MAIT cells in the lung parenchyma, we used human lungs not suitable for transplant as an essential source of lung-derived immune cells through a collaboration with the Pacific Northwest Transplant Bank in Portland, Oregon (**Supplemental Fig. 1**). Unfortunately, matched peripheral blood was not available from these donors. We used flow cytometry and a MR1-tetramer bound with either an activating ligand, 5-(2-oxopropylideneamino)-6-D-ribitylaminouracil, 5-OP-RU (MR1/5-OP-RU), or a non-activating ligand, 6-formyl pterin, 6-FP (MR1/6-FP). We quantified MAIT cells as a proportion of all live CD3^+^ T cells that stained with MR1/5-OP-RU above the minimal background staining with MR1/6-FP (**Fig. 1A**). MAIT cells comprised a higher proportion of all CD3^+^ T cells in the lung (median 2.15%, range 0.37-4.90%) compared to those from PBMC (median 0.93%, range 0.27-1.29%) (p=0.004) (**Fig. 1B**). In contrast to MAIT cells from PBMC, roughly half of the MAIT cells in the lung did not express high levels of the co-receptors CD8 or CD4 (CD8^-^CD4^-^) (median 57%) (**Fig. 1A, C**). Figure 1A shows two examples of co-receptor expression by MAIT cells. This pattern of CD8 or CD4 expression was also mirrored by non-MAIT lung derived CD3^+^ T cells (**Fig. 1C**). As the proportion of CD8-CD4-cells was higher than expected, we postulated that this diminished co-receptor expression was the result of the enzymatic processing of the lung tissue. However, we were readily able to discern these populations in the lung by flow cytometry (**Supplemental Fig. 2**). MAIT cells in the lung, as has been established for MAIT cells in the blood, are of the effector memory T cell compartment (**Fig. 1D** and **E**). Similarly, the majority of CD4-T cells within matched lung donors were also within the effector memory T cell compartment. On average, as a subset of CD3^+^ T cells, MR1-tetramer^+^ MAIT cells are 2.3-fold more abundant in the lung than in the blood, show heterogeneity based on co-receptor expression, and exclusively display an effector memory T cell phenotype.

**Figure 1.**
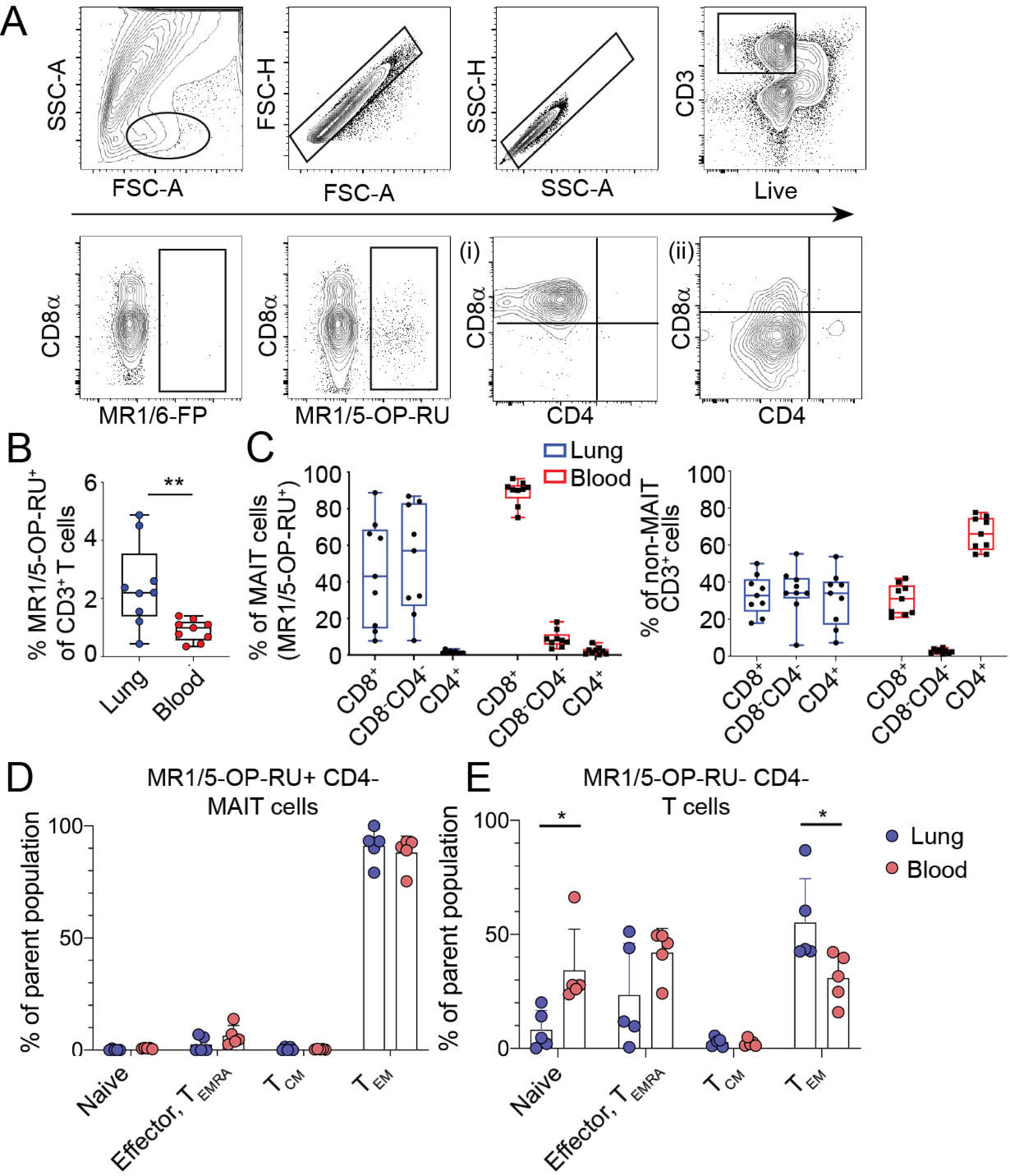
MR1-tetramer^+^ MAIT cells are an abundant T cell subset in the human lung. Lung-derived cells were stained with antibodies to T cell identification markers and the MR1-tetramer displaying either the activating 5-OP-RU ligand or the inert 6-FP ligand. A. MAIT cells were identified by the gating strategy shown on an example lung tissue as CD3^+^ MR1/5-OP-RU^+^. Staining with MR1 tetramer/6-FP was used as a negative control. MR1/5-OP-RU^+^ cells are plotted by CD4 vs. CD8*α* expression where (i) shows a lung sample where the majority of MAIT cells express CD8*α* while (ii) shows a sample where MAIT cells were mostly CD8^-^CD4^-^. B. Frequencies of MR1/5-OP-RU^+^ cells of total CD3^+^ T cells in lung parenchyma compared to the PBMC are shown by box plot. Horizontal bars indicate the median values. Statistical significance of the difference between groups was determined using the nonparametric Mann–Whitney U-test. ** *P* < 0.01, n = 9 per tissue, age range of lung donors 6-64 years.C. MR1/5-OP-RU^+^ MAIT cells (left) or non-MAIT T cells (right) were stained for their expression of CD4 and CD8 co-receptor by flow cytometry. Cells derived from lung samples are summarized by blue box plots, PBMC samples by red box plots, n = 9 per tissue. Graphs showing the % of (D) MR1/5-OP-RU^+^ CD4-MAIT cells or (E) MR1/5-OP-RU-, CD4-T cells in the lung and PBMC within common CD8^+^ T cell memory compartments. T cells were identified as naïve (CD45RO^-^, CD62L^+^, CCR7^+^), effector or TEMRA (CD45RO^-^, CD45RA^+^ or (-) CD62L^-^, CCR7^-^), central memory (CD45RO^+^, CD62L^+^, CCR7^+^), and effector memory (CD45RO^+^, CD62L^-^, CCR7^-^). Age range of lung donors: 22-51 years \**P* < 0.05, n = 5 per tissue, SD shown.

**Figure 2.**
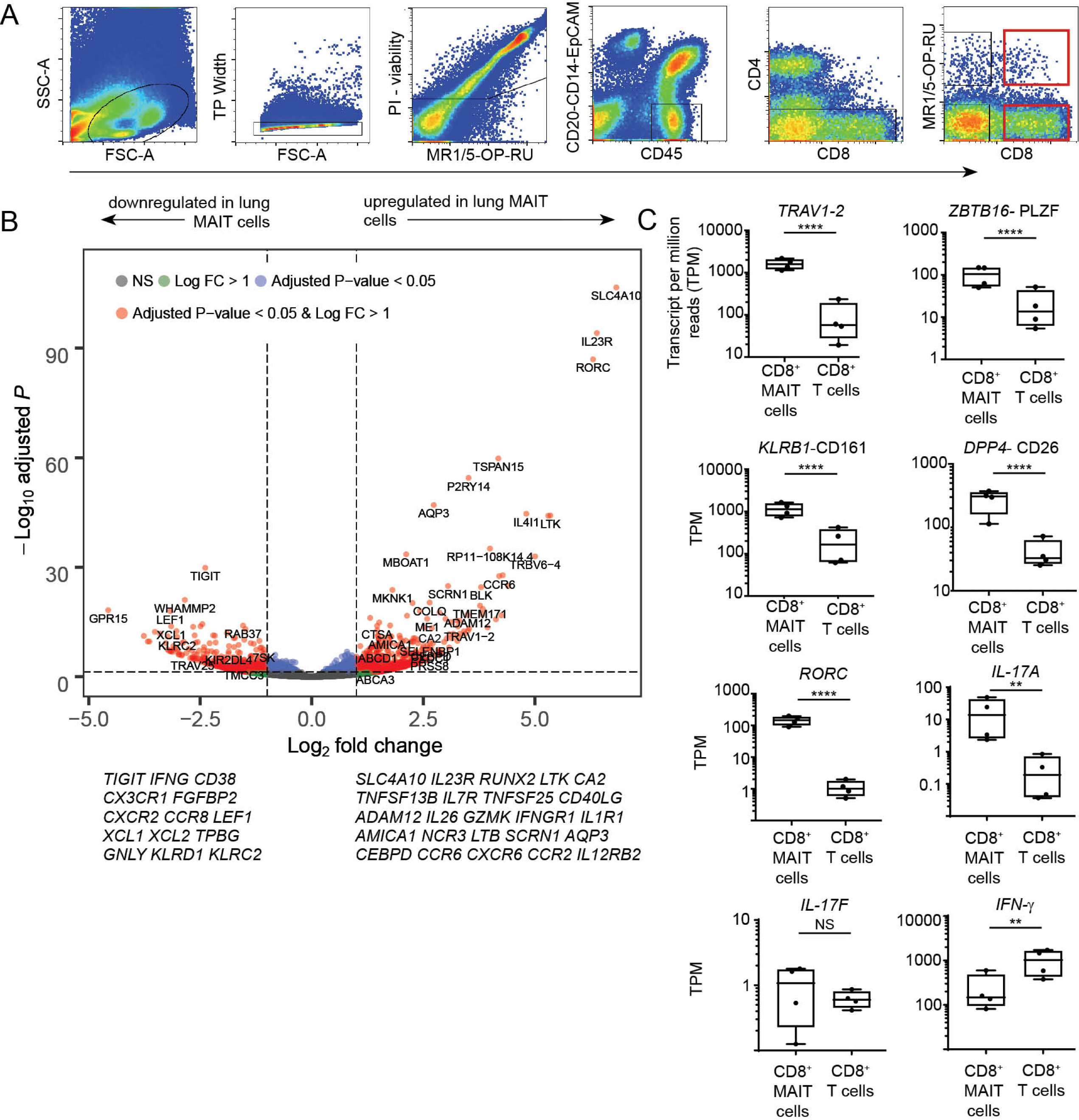
RNA-seq profiling of MAIT cell and CD8^+^ T cell subsets in human lungs. RNA-sequencing was performed on T cells from four lung samples and five PBMC samples that were FACS-sorted using the gating strategy shown in (A). Sorted MAIT cells were PI-/CD45+/CD20-CD14-EpCAM-/CD4- and MR1/5-OP-RU+ and CD8+. Non-MAIT cells were PI-/CD45+/CD20-CD14-EpCAM-/CD4- and CD8^+^. A representative FACS plot from a lung sample is shown. Age range of lung donors: 11-67 years. B. Volcano plot depicting differentially expressed genes of lung-derived CD8^+^ MAIT cells compared to non-MAIT CD8^+^ T cells. Significantly differentially expressed genes (log2 fold change >1 or <-1, and FDR *P*-value *≤* 0.05) are represented as red. Less significantly different genes are represented as blue, green or gray depending upon the significance thresholds that they fell below. C. Box plots of transcript per million (TPM) reads of genes commonly associated with MAIT cells, from RNA-seq samples of CD8^+^ MAIT cells or non-MAIT CD8^+^ T cells derived from four lung samples. Box plots indicate the median and range of the TPM RNA-seq reads. FDR \**P* < 0.05, \*\**P* < 0.01, \*\*\*\**P* < 0.0001.

### RNA-sequencing of human lung-derived MAIT cells

To determine if MAIT cells from the lung display a profile distinct from non-MAIT CD8^+^ T cells, we performed differential gene expression analysis on CD8^+^ MAIT cells and CD8^+^ non-MAIT cells isolated from the same lung. MAIT cells were identified by staining with the MR1/5-OP-RU tetramer and expression of CD8 (**Fig. 2A** red gates). Non-MAIT CD8^+^ T cells were defined by absence of staining with the MR1/5-OP-RU tetramer and expression of CD8. Matched MAIT and non-MAIT cells were collected from four different donors and RNA-sequencing was performed. While we also collected, and performed gene expression analysis on both the CD8^+^ and the CD8-CD4-, tetramer positive cells, we have focused on the CD8^+^ cells as we did not find major differences in these populations using principal component analysis. Additionally, we found that sorting based on the expression of the co-receptor enhanced the quality of the gene-expression analysis in that we did not observe evidence for B cell contamination that while rare, might have been the result of non-specific tetramer binding. Gene signatures of human lung resident CD8^+^ T cells have recently been published ^28, 29^. We observed that the MAIT and non-MAIT cells derived from lung tissue expressed many tissue residency-associated transcripts, such as CD69 and CD103 (*ITGAE*), supporting that our cells were derived from the lung tissue not vasculature (**Supplemental Fig. 3**).

**Figure 3.**
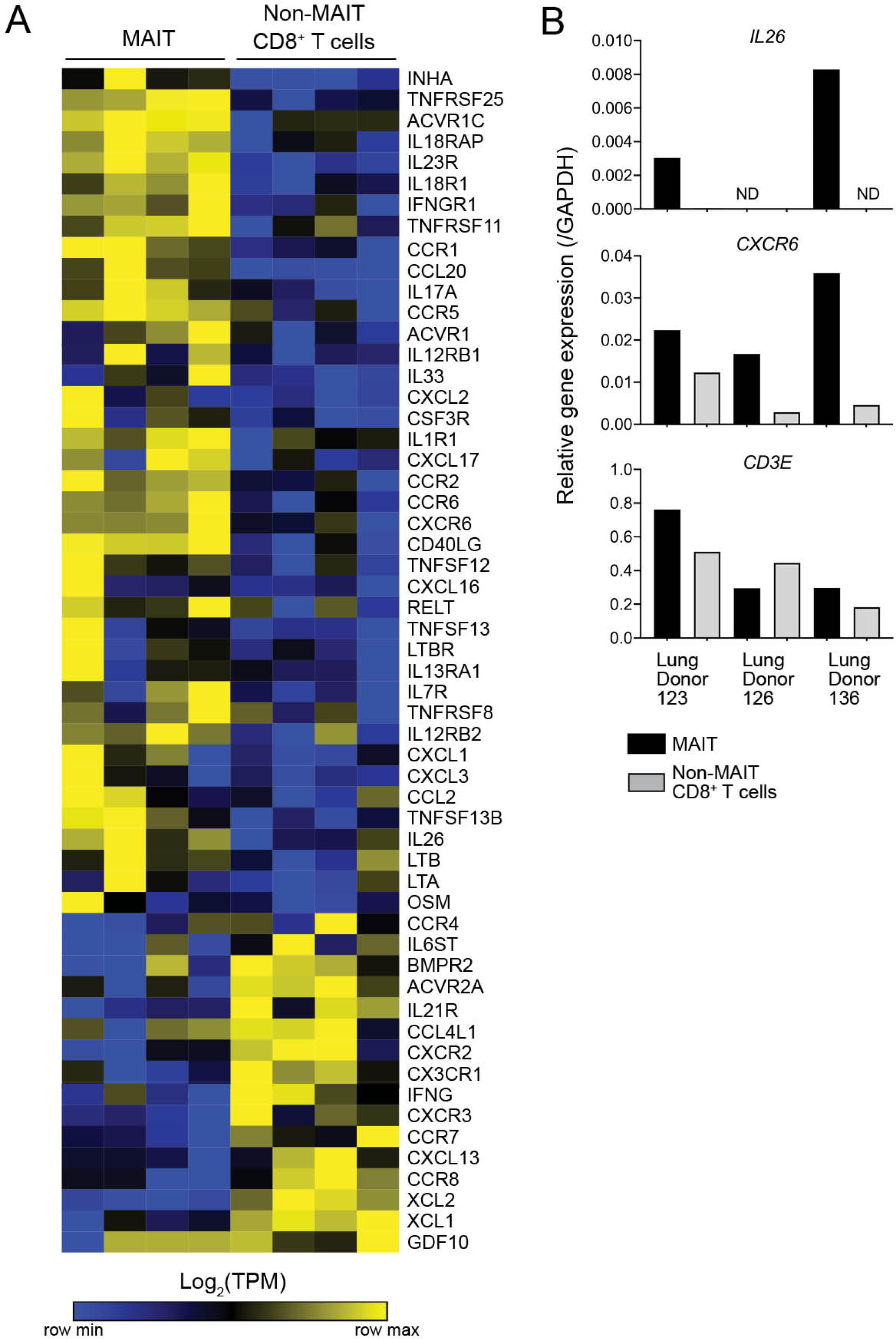
Selective expression of cytokine and cytokine receptor genes by MAIT cells in the lung. A. A heatmap showing transcripts per million reads (TPM) of cytokine and cytokine receptor (KEGG: hsa04060) gene expression in CD8^+^ MAIT cells and CD8^+^ non-MAIT cells (matched) from four lung donors, normalized by row (key). Genes selected from KEGG pathway were differentially expressed by MAIT or non-MAIT CD8^+^ T cells (FDR *≤* 0.05) and had an average TPM > 10 in at least one of the two subsets of cells. Genes and samples ordered by unsupervised hierarchical clustering using one minus spearman rank correlation. B. Bar graphs displaying quantitative PCR relative gene expression of *IL26, CXCR6,* and *CD3E*, normalized to each sample’s expression of *GAPDH*. Samples comprised of MAIT cells or non-MAIT CD8+ T cells were FACS-sorted from three additional lung donors from those in panel (A). Samples were tested in triplicate and averages are displayed. Age range of lung donors: 22-51 years.

### The distinct transcriptome of lung-derived MAIT cells as compared to non-MAIT CD8^+^ T cells

Differential gene expression analysis of the lung samples showed a large number of genes significantly (FDR-adjusted p-value *≤* 0.05) expressed in MAIT cells as compared to non-MAIT CD8^+^ T cells (**Fig. 2B and Supplemental Table 1**). Many upregulated genes are in line with transcriptional profiles of MAIT cells in the blood, suggesting an underlying homogenous program in MAIT cell populations, extending to those residing in the human lung ^30–33^ (**Fig. 2B and Supplemental Table 1**). Significantly expressed genes identified within MAIT cells in the lung included genes associated with MAIT cell identity and function, such as the invariant TCR *TRAV1-2*, the transcription factors *ZBTB16* (PLZF) and *RORC*, *DPP4* (CD26), *KLRB1* (CD161) (**Fig. 2C**) and *IL23R*. In addition to RORC, we found robust expression of *IL17A* (**Fig. 2C**), concordant with our observations in both peripheral blood ^34^ and the lung ^15^ . The highest significantly upregulated gene in lung MAIT cells was *SLC4A10. SLC4A10* encodes a sodium bicarbonate transporter NCBE, possibly associated with pH homeostasis. Similarly, we found preferential expression of *CA2*, a gene associated with pH homeostasis, but whose function in T cells is not known. Genes associated with long term survival of T cells including *RUNX2*, which plays a T cell intrinsic role in the long-term persistence of memory CD8^+^ T cells after LCMV infection in mice^35^, *LTK, TNFSF13B, IL7R*, and *TNFSF25* were more highly expressed by MAIT cells. Specific costimulatory receptors were also enriched in MAIT cells including *CD40LG* and *ADAM12* ^36^. The MAIT cell population differentially expressed the following cytotoxicity related-genes: *IL26, GZMK, IL1RI, AMICA1, NCR3, LTB*, and *SCRN1.* Interestingly, MAIT cells in the lung expressed high levels of transcript for cytotoxic molecule granzyme B which contrast with the low granzyme B expression that has been reported in circulating MAIT cells ^23^. Finally, genes involved in cell extravasation and recruitment to inflamed tissue including *CEBPD* ^37^ (CCAAT/enhancer-binding protein delta)*, CCR6, AQP3, CXCR6, CCR2, and CCR1* were more highly expressed in lung MAIT cells.

We also found genes that were significantly less expressed in the MAIT cell population compared to the non-MAIT lung derived CD8^+^ T cells. Some of these genes are associated with T cell activation after antigen experience, and include, *IFNG, CD38, CX3CR1, TIGIT, FGFBP2, CXCR2*, and *CCR8* (**Fig. 2B and Supplemental Table 1**). *LEF1*, a transcription factor in the WNT/*β*-catenin pathway, was also significantly downregulated in lung MAIT cells compared to lung non-MAIT cells. Moreover, an inhibitor of the WNT/*β*-catenin pathway was more highly expressed in lung MAIT cells, *TPBG*. Lastly, MAIT cells expressed less of the genes encoding the innate receptors CD94 and NKG2C (*KLRD1, KLRC2*) than non-MAIT CD8^+^ T cells, genes associated with TCR independent activation of MAIT cells in the blood ^19^.

Upon pathway analysis of the lung MAIT cell associated genes, we found them to be enriched with the Cytokine-Cytokine receptor interaction (hsa04060) KEGG pathway (**Fig. 3A**); thus suggesting unique pathways of sensing the lung microenvironment by MAIT cells. As shown in Figure 3, unsupervised clustering reveals subsets of cytokines, chemokines, and their respective receptors that are significantly differentially expressed within lung MAIT cells compared to non-MAIT CD8^+^ T cells. We validated two of these gene patterns with qPCR using an additional cohort of three lung organ donors and the gene *CD3E* as a positive control. For the *IL26* gene, transcript was found in MAIT cells in two of three donors and not in non-MAIT CD8^+^ T cells (**Fig. 3B**). For *CXCR6*, encoding a receptor which can integrate epidermal innate immune chemokine signals, we observed higher transcript in all three additional lung MAIT cell samples (**Fig. 3B**). These data help to delineate cytokine signaling pathways that MAIT cells could use for environmental surveillance as well as modulation of local inflammation at lung barrier tissue. Collectively, genes differentially expressed by lung MAIT cells are associated with amplifying pro-inflammatory signals and T cell chemotactic ligands.

### CD8^+^ and CD8^-^CD4^-^ MAIT cells derived from the lung share a similar transcriptomic profile

As described above, we have been able to derive a core set of genes gene associated with lung-resident CD8^+^ MAIT cells. As CD8^-^CD4^-^ MAIT cells are common in the lung (**Fig. 1**), we wanted to understand whether they shared the same core gene set as CD8^+^ MAIT cells. A study by Dias et al. found that blood-derived CD8^-^CD4^-^ MAIT cells were functionally distinct from MAIT cells expressing CD8 ^32^ and that CD8^-^CD4^-^ MAIT cells reflect an activated pool of CD8^+^ MAIT cells. Using qPCR, we confirmed that CD8 gene expression was present uniquely on the CD8^+^ subset (**Supplemental Fig. 4**). To independently derive a gene set for CD8^-^CD4^-^ MAIT cells, we analyzed differential gene expression within the transcriptomes of lung CD8^-^CD4^-^ MAIT and CD8^-^CD4^-^ non-MAIT cells from four lung donors. Here, we defined genes significantly (FDR *≤* 0.05) expressed in CD8^-^CD4^-^ MAIT cells as compared to CD8^-^CD4^-^ non-MAIT cells (**Supplemental Fig. 4**). This analysis revealed a highly similar gene expression pattern to the one derived for CD8^+^ MAIT cells (**Fig. 2B**). As we did not observe differences in the activation status, we postulate that these populations reflect similar transient activation histories in the lung. These data also support our earlier comparative emphasis on CD8^+^ cells.

**Figure 4.**
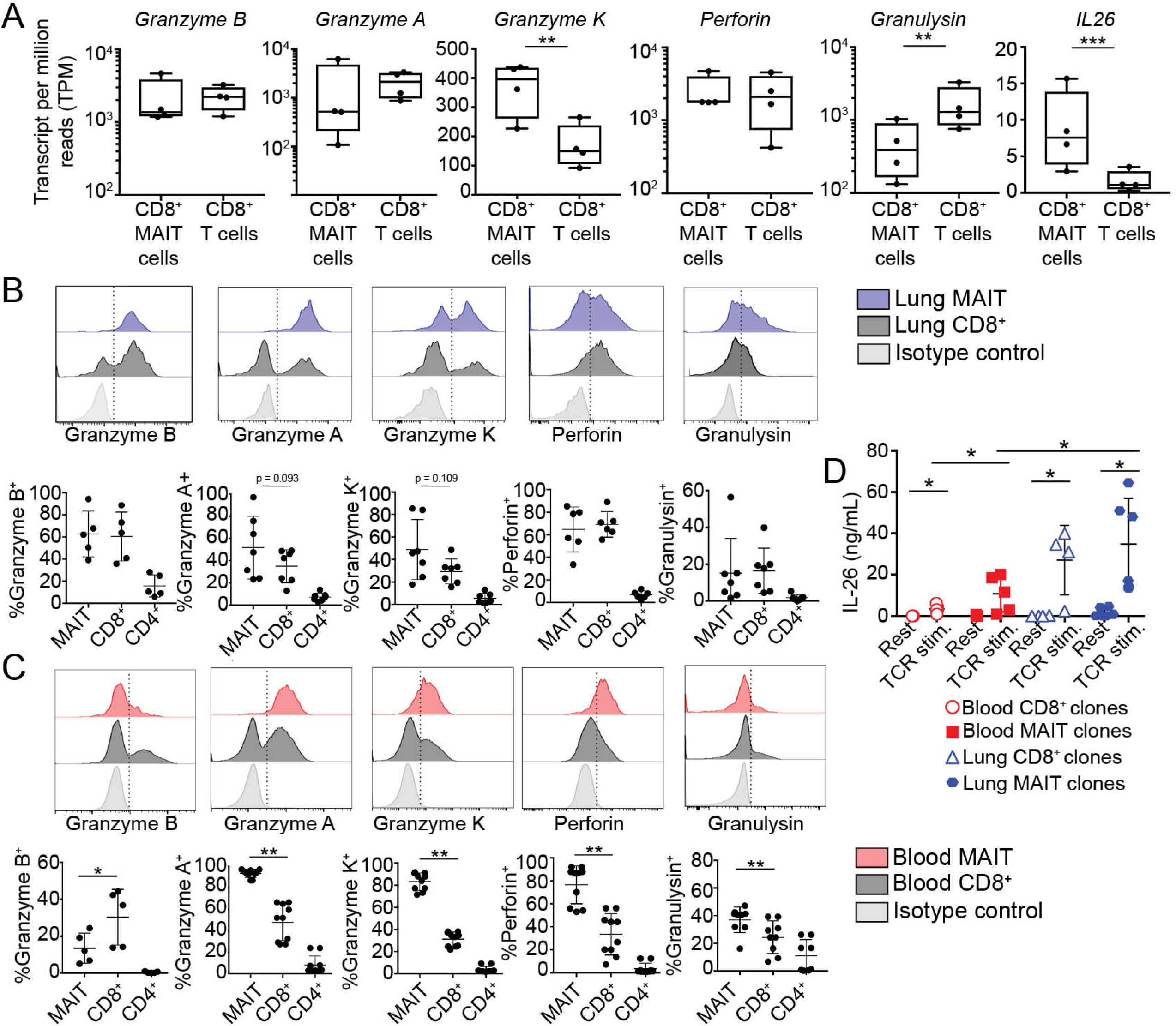
MAIT cells in the lung are enriched for preformed cytotoxic molecules and can produce IL-26. A. Transcripts per million (TPM) reads of cytotoxic T cell function related genes from paired lung RNA-seq samples. B. Flow cytometry on lung samples to measure intracellular levels of pre-formed cytotoxic molecules in MAIT cells (blue) or non-MAIT CD8^+^ T cells (gray). Top chart is a representative example of the staining and gating. Summary of all samples is the bottom chart. N = 5-7 biological replicates; age range of donors is 6 to 69 years. C. Flow cytometry on PBMC samples to measure intracellular levels of pre-formed cytotoxic molecules in MAIT cells (red) or non-MAIT CD8^+^ T cells (gray). Top chart is a representative example of the staining and gating. Summary of all samples is the bottom chart. n = 5-10 biological replicates. D. T cell clones derived from PBMC or lung BAL fluid were rested or stimulated with T-cell activation beads (anti-CD3/28) overnight and then the collected supernatant was tested by ELISA for the presence of IL-26 cytokine. Assay was performed twice with the same results and one of the experiments is displayed. Nonparametric T tests were used to test differences between groups. *P*≤*0.05. **p*≤*0.01

### MAIT cells in the lung have a polycytotoxic phenotype and secrete IL-26

Cytolytic properties of immune cells, including MAIT cells, are associated with the elimination of intracellular bacterial infection, either through the induction of apoptosis of the target cells, or via the introduction of anti-bacterial proteins. Traditionally in humans, cytotoxic CD8^+^ and CD4^+^ T cells contain granzyme B, perforin, and granulysin in preformed granules, allowing for rapid delivery to the target cell ^38–41^. MAIT cells also express cytotoxic molecules and are capable of lysing bacterially infected cells ranging from macrophages to epithelial cells. MAIT cell degranulation is associated with MR1 antigen presentation ^24^. Circulating MAIT cells express less granzyme B compared to non-MAIT T cells ^23^. They must be licensed to produce granzyme B in the context of infection, TCR or MR1-independent cytokine stimulation. Proteomics analysis of blood-derived MAIT cells identified multiple cytotoxicity-related proteins with pronounced expression in MAIT cells, compared to non-MAIT CD8^+^ T cells, in the context of *E. coli* infection ^42^. Correspondingly lung-derived MAIT cells expressed high levels of transcript for traditional cytolytic proteins including granzyme B, granzyme A, granzyme K, perforin, and granulysin (**Fig. 4A, Supplemental Table 1**). Cytotoxicity related-genes specifically enriched in MAIT cells over non-MAIT CD8^+^ T cells included: *IL26, GZMK, LTB, IL1RI, AMICA1, NCR3*, and *SCRN1.* (**Supplemental Table 1**). Through populational RNA-seq analysis, we could not distinguish whether differences were at the individual cell level or the population level.

To validate the repertoire of preformed cytotoxic proteins in lung MAIT cells compared to non-MAIT cells at the single cell level, we used flow cytometry as well as ELISA of cell supernatants. We used intracellular flow cytometry on *ex vivo* lung samples to measure expression of proteins corresponding to the genes listed in Fig. 4A at the single cell level. In concordance with the gene expression patterns, a high proportion of lung-derived MAIT cells and CD8^+^ non-MAIT cells expressed pre-formed cytotoxic proteins. We observed a trend of more MAIT cells expressing granzyme A and K compared to non-MAIT CD8^+^ lung T cells. Specifically, on average, 62%, 52%, 50%, 63%, and 17% of MAIT cells expressed preformed granzyme B, granzyme A, granzyme K, perforin, and granulysin in the lung, respectively (**Fig. 4B**). In contrast, 60%, 34%, 30%, 69%, and 18% of non-MAIT CD8^+^ T cells in the lung expressed granzyme B, granzyme A, granzyme K, perforin, and granulysin, respectively. A minority of CD4^+^ T cells in the lung expressed cytotoxic proteins. To compare the levels of cytotoxic proteins between MAIT cells derived from lung and blood, we next assessed the intracellular levels of the same proteins from T cell populations in PBMC donors (**Fig. 4C**). While a significantly higher proportion of MAIT cells from the blood express granzyme A, granzyme K, perforin, and granulysin than non-MAIT CD8^+^ T cells, they also expressed less granzyme B than non-MAIT CD8^+^ T cells and lung-derived MAIT cells (comparison not shown; p value = 0.008). Overall, we found more heterogeneity in the frequency of each cell type’s expression of cytotoxic proteins in the lung, which could indicate infection history, priming environment, or recent exposure to activating ligand in the lung. Polycytotoxicity has been defined as the co-expression of perforin, granulysin, and granzyme ^43^. Therefore, MAIT cells represent a polycytotoxic T cell population in the lung, and in contrast to MAIT cells in the blood, express preformed granzyme B.

Lung-derived MAIT cells expressed significant higher levels of *IL26* (**Fig. 3B, 4A**). IL-26 is an IL-20 family cytokine that signals via a heterodimeric receptor expressed on epithelial cells ^44, 45^. It has both signaling characteristics and cationic properties that allow it to bind to DNA and also disrupt bacterial membranes^46^. Interesting, IL-26 has direct antimicrobial activity against mycobacteria, including *Mycobacterium tuberculosis* ^47^. Transcriptomic studies of blood derived MAIT cells show presence of *IL26* upon stimulation through the TCR or with IL-12/18 ^48 31, 49^. We used ELISA to determine the presence of IL-26 from a panel of both MAIT and non-MAIT T cell clones derived from either the blood or bronchioalveolar lavage (**Fig. 4D**). TCR stimulation was performed using anti-CD3/28 coated beads. We observed that all of the T cell clones were able to make IL-26 when stimulated through the TCR (**Fig. 4D**) although MAIT T cell clones derived from the lung made more IL-26 than those derived from the blood. Taken together, MAIT cells in the lung exhibit a polycytotoxic phenotype as a high proportion of them express several granzymes, perforin, granulysin, and can produce IL-26.

### MAIT cells express distinct cytokine receptors including IL-12 and IFN-γ receptors

MR1-independent cytokine-driven responses to bacterial infection have been observed from circulating and liver-derived MAIT cells. Specifically, these responses have been linked to IL-12 and IL-18. Furthermore, IL-18 and IL-12 p40 are important for the *in vivo* antibacterial activity of MAIT cells in multiple disease models ^1, 2, 4^. Transcriptome analysis revealed higher gene expression of specific cytokine receptors (**Fig. 3A**), including *IL23R*, *IFNGR1* (**Fig. 5A**), and *IL12RB1/2* (**Fig. 5B, C)** by lung MAIT cells as compared to non-MAIT cells. To assess whether this transcriptional pattern was matched at the protein level, we measured cytokine receptors by flow cytometry on cell populations in blood and lung samples. Here, we observed that a significantly higher proportion of lung and blood MAIT cells expressed IFN-*γ* receptor compared to non-MAIT CD8^+^ T cells (**Fig. 5A**). The IL-12 receptor is comprised of a heterodimer of IL-12 receptor *β*1 and IL-12 receptor *β*2. Given that the IL-12 receptor heterodimer is upregulated during inflammation, we measured each receptor separately on resting cells or following four days of stimulation with anti-CD3/28, IL-2, and IL-12. Due to our inability to obtain sufficient lung MAIT cells to culture, these experiments were performed with blood-derived cells. Here, we observed that while a minor fraction of MAIT cells and non-MAIT CD8^+^ T cells expressed IL-12R*β*1 at rest, nearly all of the cells expressed IL-12 receptor *β*1 following stimulation (**Fig. 5B**). Additionally, MAIT cells expressed more of the receptor as suggested by comparing the average mean fluorescence intensities of the stimulated populations (CD8^+^: 404 vs. MAIT: 1096). As IL-12 receptor *β*1 can pair with IL-23 receptor (for IL-23) or IL-12 receptor *β*2 (for IL-12), we also measured IL-23 receptor levels on resting cells and after stimulation. We did not observe IL-23 receptor expression from any of the lung or PBMC samples by flow cytometry using multiple available antibodies (data not shown). However, when we measured IL-12 receptor *β*2 under the same stimulation conditions, we observed that a proportion (on average 19%) of MAIT cells expressed this receptor (**Fig. 5C**). IL-12 receptor *β*2 was not expressed by stimulated non-MAIT CD8^+^ T cells.

**Figure 5.**
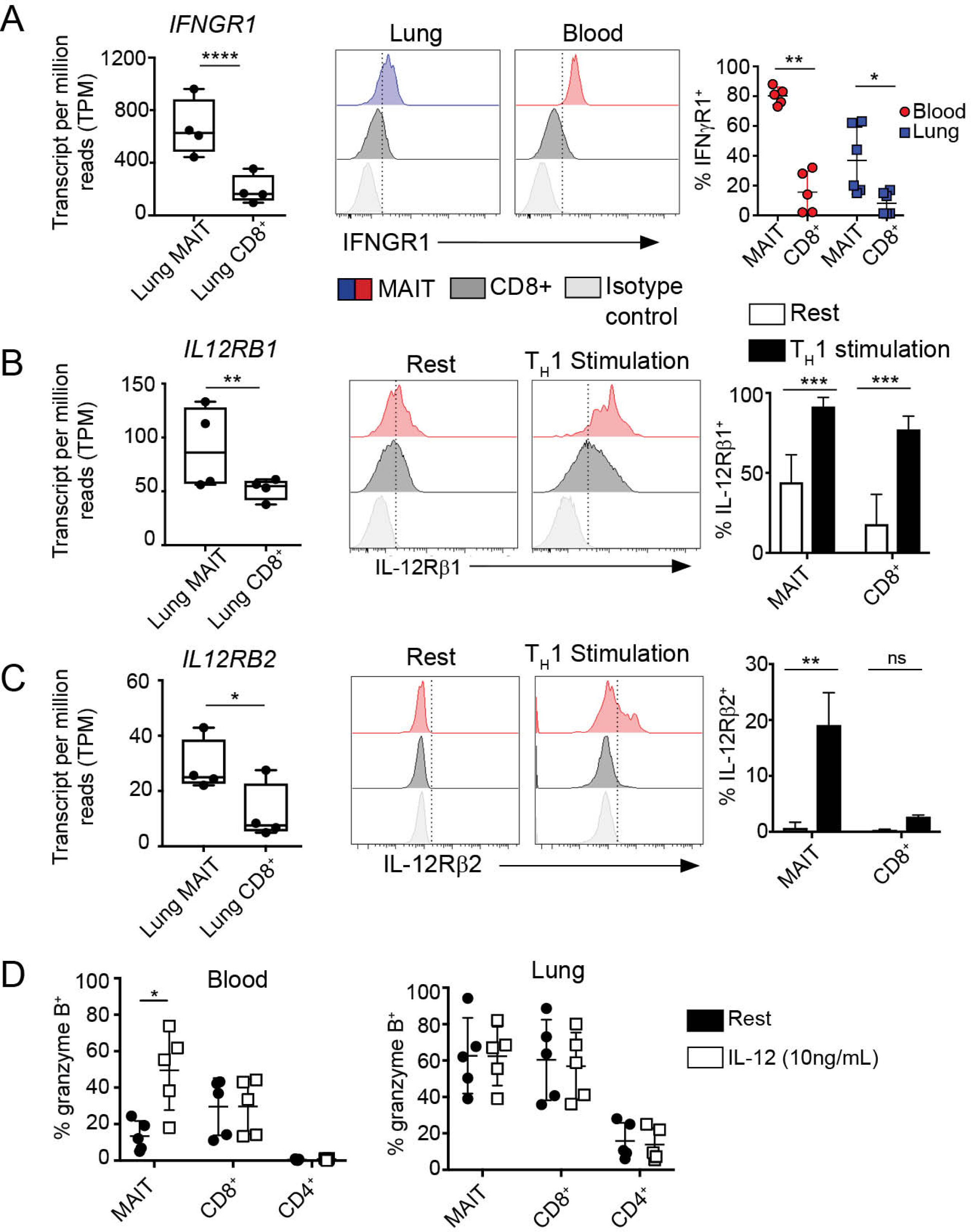
IL-12 upregulates granzyme B production in circulating MAIT cells. A. Transcripts per million (TPM) of interferon-gamma receptor *(IFNGR1*) in lung MAIT and lung non-MAIT CD8^+^ T cells. *Ex vivo* lung and blood samples were stained with antibody specific for IFN-*γ* receptor on the surface. Histograms show an example of flow cytometry quantification of IFN-*γ* receptor on the gated populations, MR1/5-OP-RU^+^ MAIT cells, MR1/5-OP-RU(-) non MAIT CD8^+^ T cells, and isotype control staining of CD8^+^ T cells. A summary of the frequency of IFNGR1^+^ cells is shown on the right, n = 5. The % of the gated population staining above the isotype control is plotted on the y-axis. ; age range of donors is 6 to 69 years. B, C. TPM of interleukin 12 receptor *β*1 (B) or *β*2 (C) in lung MAIT, lung non-MAIT CD8^+^ T cells samples. Validation at the protein level was performed on PBMC that were rested or stimulated for four days under Th1 conditions including anti-CD3/28, IL-2, and IL12. Lymphocytes were then stained for surface IL-12R*β*1 (B) or IL-12R*β*2 (C). Staining of MAIT cells (red), non-MAIT CD8^+^ T cells (black) or the isotype control (gray) are shown by histograms. Summary of the frequency of IL-12R*β*1 or *β*2 expression by MR1/5-OP-RU^+^ MAIT cells or non MAIT CD8^+^ T cells. Bar graphs of IL-12R*β*1 expression represent the mean and SD of the frequency of positive cells within that gated population in 5 biological replicates repeated three times. Bar graphs of IL-12R*β*2 expression represent the mean and SD of two technical replicates. D. Cells derived from PBMC or lung samples were either rested (black circles) overnight or in the presence of IL-12 cytokine (white circles, 10 ng/mL) and then stained intracellularly for granzyme B. The frequency of granzyme B positive cells is plotted from 5 PBMC donors (top) and 5 lung donors (bottom). Representative results of three independent experiments are shown, n = 5 biological replicates, age range of donors is 6 to 60 years. Nonparametric T tests were used to test differences, *p*≤*0.05, **p*≤*0.01

### MAIT cells respond to IL-12 by upregulating intracellular granzyme B

Given that few blood-derived MAIT cells expressed granzyme B (**Fig. 4C**), MAIT cells uniquely expressed IFN-*γ* and IL-12 receptors, and that these cytokines have been implicated in control of intracellular infection, we assessed whether IFN-*γ* and IL-12 could regulate cytotoxic capacity of MAIT cells. To do this, we tested whether IFN-*γ* receptor-, IL-12 receptor-, or TCR-stimulation would regulate the intracellular levels of preformed cytotoxic proteins in MAIT cells or non-MAIT T cells derived from PBMC (**Fig. 5D** top) and the lung (**Fig. 5D** bottom). We assessed the frequency of each cell type that expressed granzyme B, granzyme A, granzyme K, perforin, or granulysin by flow cytometry before and after 18-hour stimulation. We observed in blood MAIT cells that only the addition of IL-12 upregulated granzyme B but no other cytotoxic proteins (**Fig. 5D**, **Supplemental Fig. 5**). We did not observe changes with IFN-*γ* or TCR stimulation in our *in vitro* assay (**Supplemental Fig. 6**). The functional signaling consequences of the highly expressed IFN-*γ* receptor on MAIT cells remains to be determined. We note that granzyme B expression is inducible with IL-12 cytokine and thus, IL-12 can license the cytotoxic capacity of circulating MAIT cells which we hypothesize contributes to the higher frequency of granzyme B^+^ MAIT cells in the lung.

## Discussion

MAIT cells are an abundant human T cell subset that shares qualities of the innate and adaptive immune system through their recognition of small microbial metabolites presented by MR1. These cells are of particular interest in the setting of respiratory bacterial infection due to their abundance in the airway and demonstrated ability to respond to cells infected with bacteria ^20, 50, 51^. However, an in-depth comparative analysis to reveal fundamental biological pathways of MAIT cells in the human lung has not yet been performed. We undertook a transcriptomic analysis of MAIT cells compared to non-MAIT CD8^+^ T cells in the human lung. We found that MAIT cells in the lung display a polycytotoxic phenotype and sense their environment through selective cytokine responsiveness. The lung-derived MAIT cell’s gene expression aligns closely with transcriptomics of blood-derived MAIT cells and other innate-like T cell compartments.

Cytokine receptors allow cells to respond to their microenvironment. We found that lung MAIT cells expressed a unique set of cytokine receptors compared to non-MAIT CD8^+^ T cells which would contribute to regulating their functions in the lung. The IL-12 receptor and IFN-*γ* receptor were expressed at the protein level by MAIT cells as well. Our staining of IL-12 receptor on MAIT cells supports the recent finding of its expression on MAIT cells derived from lung explants ^27^. MAIT cells can become activated by IL-12 and IL-18 in humans, a property thought to be distinctly linked to cell populations expressing *KLRB1*/CD161 ^52^. A role for IL-12 activation of MAIT cells, independent from MR1, has been described in *in vivo* models of bacterial respiratory infections ^2^. As IL-12 is made early in infection by macrophages, DCs, and neutrophils, our data supports the hypothesis that IL-12 receptor expression in MAIT cells facilitates the initial response to infection in the lung.

We observed that lung MAIT cells distinctively expressed transcript for IFN-*γ* receptor compared to non-MAIT CD8^+^ T cells. IFN-*γ* cytokine is made upon activation of subsets of NK and T cells and is a critical cytokine in defense against intracellular microbial infection ^53^. Given the known roles of signaling through this cytokine, IFN-*γ* receptor activity in MAIT cells likely helps in their polarization towards a Th1 and/or Th17 phenotype and helps them to amplify local cellular immune activation ^54^. We also note the expression of the transcription factor *RORC*, known to be associated with the expression of the cytokine IL-17 ^55^. While we did observe the expression of *IL17A*, we have not observed protein expression in either peripheral blood MAITs ^34^ or those in the lung ^14^. Interestingly, Cole et al ^56^ have recently found that MAIT cells express IL-17F rather than IL-17A. However, we did not find expression of IL-17F.

T cells can directly mediate antimicrobial responses through the release of antimicrobial proteins. Here, we confirmed the expression in MAITs of *GZMK, PRF* and *GNLY*, characteristic of NK cells and CD8^+^ polycytotoxic T lymphocytes ^39^ and *IL26*, characteristic of Th17 cells ^46^, regardless of tissue distribution, which we confirmed by either flow cytometry or ELISA. An analysis of immune-related genes expressed in T cells from human intestinal biopsies also linked *GZMK* to MAIT cells ^22^. Together, granzyme B, perforin and granulysin act synergistically to kill intracellular bacteria ^38, 43, 57^. Perforin disrupts the target eukaryotic cell membrane, allowing other cytotoxic proteins to gain access to the inside of the cell. Granzymes are proteases with many host and microbial molecular targets. For example, granzymes can activate host caspases, deactivate host oxidative stress defense proteins, or deactivate bacterial metabolic pathways ^57, 58^. Furthermore, each granzyme has different proteolytic targets. As MAIT cells expressed high levels of granzyme B, A, and K, this suggests a broad proteolytic spectrum. Granulysin can bind to bacteria damaging its membrane or cell wall ^38^. Unexpectedly, the gene encoding the antimicrobial protein, IL-26, was significantly upregulated in MAIT cells. Notably, both MAIT cell clones and non-MAIT cell clones derived from the lung produced IL-26 which required TCR-stimulation. This discrepancy from the core RNA signature pattern could be explained if MAIT cells in the lung are licensed to make IL-26. IL-26 kills extracellular bacteria ^46^ but can enter macrophages and exert an antimicrobial activity against intracellular bacteria ^59^. This suggests that MAIT cells are armed to have broad antimicrobial effector function.

We hypothesized that this polycytotoxic activity could be enhanced as a result of their unique cytokine receptors. In contrast to blood MAIT cells, which express little granzyme B, we found that lung MAIT cells contain preformed granzyme B. Furthermore, we found that *in vitro* stimulation of circulating MAIT cells, but not non-MAIT CD8^+^ T cells, with IL-12, but not IFN-*γ*, upregulated intracellular levels of granzyme B. This result would suggest that a cytokine rich lung environment could “license” or augment the cytolytic activity of MAIT cells.

We acknowledge that our bulk RNA-sequencing method may have overlooked functionally distinct MAIT cell subsets. Our transcriptomic analysis was performed directly *ex vivo* so that we could examine their natural state in the lung. Moreover, we had hypothesized that the higher proportion of CD8^-^CD4^-^ MAIT cells in the lung may represent a distinct subset at the transcriptomic level and we therefore, performed RNA-seq analysis on these cells separately. Instead, we found very few significant differences in genes upregulated in CD8^-^CD4^-^ MAIT cells compared to CD8^+^ lung derived MAIT cells. Whether this reflects a common lung environment, or a common lung MAIT precursor the result of imprinting early in life from the microbiome^6^, remains to be determined.

Given MAIT cells’ positioning near the epithelium^14^, our findings raise the intriguing possibility that lung-resident MAIT cells could play a specialized role as environmental sensors and immediate effectors upon infection in the airway. Specifically, this could be the result of the release of pro-inflammatory cytokines and chemokines, which assist in the conditioning and migration of myeloid subsets ^60^, as well as by their direct anti-microbial activity. MAIT cells are also in a position to directly kill microbes through polycytotoxic mechanisms, a function which is regulated through antigen specific and innate activation. In the future, how these immune defense pathways change when they become fully activated during a microbial encounter will need to be studied in parallel. Whether these immune defense pathways at the mucosa are successful upon encounter or not could reveal key correlates of a specific immune response.

Finally, the gene profile we have observed would also suggest that MAITs in the lung are poised to tolerate a relatively harsh environment of the airway (*SLC4A10, CA2, and Aquaporin 3)*, or possibly to participate in tissue repair as has been reported by Constantinides et al., and Leng et al ^6 49^. Of relevance to human MAIT cells, Leng et al., found that following MAIT cell stimulation, preferential expression of Furin and CCL3. They then demonstrated that these genes were associated with tissue repair. While we did not observe preferential expression of these genes, we note that our cells were not specifically stimulated such that we may have missed this phenotype.

## Materials and methods

### Human subjects

All samples were collected and all experiments were conducted under protocols approved by the institutional review board at Oregon Health and Science University. PBMCs were obtained by apheresis from healthy adult donors with informed consent. De-identified lungs not suitable for transplant were obtained from the Pacific Northwest Transplant Bank (PNTB). Our exclusion criteria included significant tobacco smoking history, drowning, crushing chest injuries, lobar pneumonia, and HIV/HBV/HCV infection. Bronchoalveolar lavage fluid for generating T cell clones was obtained under a protocol approved by the UKZN Biomedical Research Ethics Committee and the Partners Institutional Review Board. The participants provided written informed consent.

### Human tissue sources of T cell populations

PBMCs were isolated from the peripheral blood of healthy donors using Ficoll-Paque gradients. Lung single cell suspensions are prepared from recently deceased donor tissue not suitable for transplant from the Pacific Northwest Transplant bank. Small cubes of lung parenchyma, devoid of airway and lymph nodes, were cut into a cold buffer of HBSS (Gibco) media supplemented with HEPES (Gibco) and PSF antibiotic (Sigma). Tissue was then digested for 30 minutes at 37 degrees C in a DMEM buffer (Gibco) supplemented with PSF antibiotics (Sigma), elastase (15 μg/mL, Worthington), trypsin I (1.5 μg/mL, Sigma), DNase I (45 μg/mL, Roche). The subsequent suspension was further dissociated using a GentleMACS dissociator (Miltenyi) using the Lung02 program. The single cell suspension is then diluted 1:1 with a buffer of HBSS (Gibco) media supplemented with 2% heat-inactivated fetal bovine serum (Gemini Bio Products), HEPES (Gibco) and PSF antibiotic (Sigma) to dilute homogenate and neutralize digest enzymes. This cell suspension is passed through successive filters in this order: metal mesh sieve filter (size 40 then 60, Sigma), and nylon cell strainer (100 μm then 40 μm, BD Falcon). The resulting cell suspension is washed in RPMI supplemented with 10% heat inactivated pooled human serum and used for experiments or cryo-preserved in heat-inactivated fetal bovine serum with 10% DMSO. Lung-derived T cell clones were isolated from bronchial alveolar lavage (BAL) samples using a protocol described in detail below.

### Flow cytometry

Cells to be analyzed for cell surface marker expression were blocked with tetramer staining buffer (PBS buffer containing 2% fetal bovine serum). PBMCs were stained with the MR1/5-OP-RU tetramer (NIH Tetramer core facility) at 0.3 nM in 25 μL volume for 45 minutes in tetramer staining buffer at room temperature in the dark. Viability and surface stains were added on top of the tetramer stain for another 20 min at 4°C in the dark. Samples were then washed in tetramer staining buffer. For intracellular staining, cells were permeabilized and fixed using the CytoFix/CytoPerm kit (BD) as directed. Antibodies to intracellular proteins were added to samples for 20 minutes at 4°C in the dark and then washed with PermWash (BD). Flow cytometry analysis was performed using a LSRFortessa flow cytometer (BD) or a CytoFLEX S flow cytometer (Beckman Coulter). Data were analyzed using FlowJo (v10.4.2). MR1/6-FP tetramer controls were used for optimal gating of MAIT cells. Doublets were excluded based on FSC-H vs. FSC-A and SSC-H vs. SSC-A; T lymphocytes were identified based on FSC-A and SSC-A and CD3 expression; dead cells were excluded based on Aqua viability dye.

| Target of Antibody | Fluorophore | Manufacturer | Clone |
| --- | --- | --- | --- |
| CD3 | PerCP Cy5.5 | BioLegend | SK7 |
| CD3 | PerCP | BD Biosciences | SK7 |
| CD3 | FITC | BioLegend | UCHT1 |
| CD3 | BV421 | BioLegend | UCHT1 |
| CD4 | BV785 | BioLegend | OKT4 |
| CD8 | APC eFluor780 | eBioscience | SK1 |
| CD8 | APC | BioLegend | HITBa |
| CD8 | FITC | BD Biosciences | RPA-T8 |
| CD14 | APC | BioLegend | HCD14 |
| TRAV1-2 | APC | Lewinsohn Lab | OF5A12 |
| TCR$\alpha\beta$ | PE | BD Biosciences | T10B9.1A-31 |
| IL-12RB1 | PE | BD Biosciences | 2.4E6 |
| Mouse IgG1 Isotype | PE | BioLegend | MOPC-21 |
| IL-12RB2 | APC | R&D Systems | 305719 |
| Mouse IgG1 Isotype | APC | BD Biosciences | MOPC-21 |
| IFNGR1 | FITC | R&D Systems | 92101 |
| Mouse IgG1 Isotype | FITC | BioLegend | MOPC-21 |
| IL-23R | FITC | R&D Systems | 218213 |
| Mouse IgG2b | FITC | BioLegend | MPC-11 |
| Granzyme A | Pac Blue | BioLegend | CB9 |
| Granzyme B | Pac Blue | BioLegend | QA16A02 |
| Granzyme K | APC | BioLegend | GM26E7 |
| Perforin | PE-Cy7 | eBioscience (Thermofisher) | dG9 |
| Granulysin | Alexa Fluor 488 | BioLegend | DH2 |
| CCR7 | PE-Cy7 | BioLegend | G043H7 |
| Ms IgG2a Isotype | PE-Cy7 | BioLegend | MOPC-173 |
| CD62L | PerCP-Cy5.5 | BioLegend | DREG-56 |
| Ms IgG1 Isotype | PerCP-Cy5.5 | BioLegend | MOPC-21 |
| CD45RO | BV421 | BioLegend | UCHL1 |
| Ms IgG2a Isotype | BV421 | BioLegend | MOPC-173 |
| CD45RA | FITC | BioLegend | HI100 |
| Ms IgG2b Isotype | FITC | BioLegend | MPC-11 |
| Mouse IgG1 Isotype | Pac Blue | BioLegend | MOPC-21 |
| Mouse IgG2b | PE-Cy7 | BioLegend | MPC-11 |

### Cell sorting of T cell populations and RNA isolation from sorted cells

Cells were surface stained using the flow cytometry staining protocol above. Specifically, cells were stained with propidium iodide viability stain, MR1/5-OP-RU tetramer, and antibodies to CD45, CD4, CD8. Lung samples were also stained with antibodies to CD20, CD14, and EpCAM for removal of B cells, myeloid cells, and epithelial cells. Live, CD45+, CD4 negative, CD14/CD20/EpCAM negative were sorted into four populations: MR1/5-OP-RU+ CD8+, MR1/5-OP-RU+ CD8 negative, MR1/5-OP-RU-negative CD8+, and MR1/5-OP-RU-negative CD8 negative. Cells were sorted into TRIzol (Thermo-Fisher) and RNA was isolated using a Direct-zol RNA-mini kit with RNA clean and concentrator columns (Zymo Research) and treated with DNase (Qiagen).

### Quantitative PCR

Lung cells were sorted directly into 500 μL of RLT buffer in 1.5 mL Eppendorf tubes. 5-OPRU tetramer negative cells were collected as 10,000 cells each and 5-OPRU tetramer positive cells from three lung donors were collected as 3489 cells, 3959 cells and 4730 cells, respectively. Total RNA was isolated using RNeasy mini kit (Qiagen) and eluted with 30 μL of nuclease free water. 9 μL of total RNA samples were proceeded to cDNA synthesis using High Capacity cDNA Reverse Transcription kit (Applied Biosystems). Then, cDNA was amplified using TaqMan PreAmp Master Mix (Applied Biosystems) with assay mix contains GAPDH (Hs02758991_g1), CD3 (Hs01062241_m1), CXCR6 (Hs00174843_m1) and IL-26 (Hs00218189_m1). The preamplification reaction was done with 14 cycles and reactions were diluted 20 times with TE buffer. The PCR reaction was prepared according to procedure of PreAmp mix and performed with StepOnePlus Real-Time PCR System (Applied Biosystems).

### RNA-Sequencing sample preparation

Sequencing libraries were constructed from total RNA using NuGEN Ovation Ultralow library kit and NuGEN Ovation RNA-seq v2 kit (NuGEN). Libraries were sequenced at OHSU Massively Parallel Sequencing Shared Resource with the Illumina HiSeq platform with single end reads and 100 base read length. An average of 65 million reads were generated for each sample. Data is deposited in NCBI SRA (BioProject ID: PRJNA830885).

### RNAseq transcriptome analysis

FastQC (https://www.bioinformatics.babraham.ac.uk/projects/fastqc/) was used to assess the quality of the raw sequence reads. Reads were then aligned to the Human genome (Hg19) using STAR (version 2.5.2b) (PMID: 23104886) allowing for a maximum of 2 mismatches per 100 bp read. Approximately 85% of reads mapped uniquely to the genome. Read counts per gene were then obtained using HTSeq (PMID: 25260700). Differential gene expression analysis was performed using DESeq2 (PMID: 25516281).

### Generation and characterization of lung derived T cell clones

Cells from BAL samples were stained with Aqua LIVE/DEAD (Invitrogen), MR1/5-OP-RU tetramer (0.3nM), *α*-CD4-FITC (clone OKT4; BioLegend), and *α*-CD8-APC-Cy7 (clone SK8; BioLegend). Live tetramer-binding cells were sorted the basis of co-receptor expression using an Influx flow cytometer (BD Biosciences), rested overnight in RPMI 1640 supplemented with 10% heat-inactivated pooled human serum and 0.5ng/ml rhIL-2, and distributed in limiting dilution format with irradiated PBMCs and irradiated B-lymphoblastoid cells in a 96-well round bottom plate. The cultures were supplemented with rhIL-2 (5ng/ml), rhIL-12 (0.5ng/ml), rhIL-7 (0.5ng/ml), rhIL-15 (0.5ng/ml) and *α*-CD3 (0.03 µg/ml). T cell clones were harvested after incubation for 20 days at 37°C.

### Expansion of T Cell Clones

T cell clones were cultured in the presence of x-rayed (3000 cGray using X-RAD320, Precision X-Ray Inc.) allogeneic PBMCs, x-rayed allogeneic LCL (6000 cGray), and anti-CD3 mAb (20 ng/ml; Orthoclone OKT3, eBioscience) in RPMI 1640 media with 10% human serum in a T-25 upright flask in a total volume of 30 ml. The cultures were supplemented with IL-2 on days 1, 4, 7, and 10 of culture. The cell cultures were washed on day 5 to remove soluble anti-CD3 mAb.

### ELISA

T cell clones were rested for 24 h in RPMI buffer containing 10% human serum and 0.5 ng/mL IL-2 Stimulated cells were mixed 1:2 with MACS Miltenyi Biotec T cell Activation beads. T cell clones were then incubated for 24 h at 37°C. Cells supernatants were collected and stored at -80°C. Cytokine secretion was quantified with an IL-26 ELISA kit (Cusabio) according to the manufacturer’s instructions.

### T cell *in vitro* stimulation assays

To measure IL-12 receptor surface expression, PBMC derived T cells were stimulated for five days at 37°C with 1μg/mL anti-CD3 and anti-CD28 and 10ng/mL of recombinant human IL-2 and recombinant human IL-12p70 (BioLegend). On day four of stimulation, cells were washed with RPMI medium with 10% human serum and 0.5 ng/mL of IL-2 before continuing incubation at 37°C. Following this five-day period, cells were stimulated with 50ng/mL of PMA and 200ng/mL of ionomycin for three hours.

### Statistical analyses

Statistical analyses were performed using GraphPad Prism 7 software (GraphPad Software Inc., San Diego, CA). Datasets were analyzed for normality using Prism. If normal, a Student’s two-tailed T test was used for comparisons in Prism. If the data were not normally distributed, the non-parametric T-test was used to assess significant differences between groups in Prism. Error bars in the figures indicate the standard deviation, standard error of the mean, or the data set range as indicated in each figure legend. *P* values *≤* 0.05 were considered significant (\**P≤*0.05; ** *P≤*0.01; *** *P≤*0.001; **** *P≤*0.0001).

### Data Availability

All relevant data are available in this manuscript and from the corresponding author.

## Acknowledgements

We thank the Pacific Northwest Transplant Bank for providing tissue samples vital to our investigation. We thank staff of both the OHSU Massively Parallel Sequencing Shared Resource and OHSU Flow Cytometry core. We thank members of the Lewinsohn lab for technical support and helpful discussions. The following reagents were obtained through the NIH Tetramer Core Facility: MR1/5-OP-RU, MR1/6FP tetramers. The MR1/5-OP-RU tetramer technology was developed jointly by J. McCluskey, J. Rossjohn, and D. Fairlie, and the material was produced by the NIH Tetramer Core Facility as permitted to be distributed by the University of Melbourne.

## Funding

This work was supported by Merit Review Awards I01BX000533 from the United States Department of Veterans Affairs Biomedical Laboratory Research and resources and the use of facilities at the VA Portland Health Care System; NIAID R01AI048090, R01AI134790, U01 AI095776, K08 AI118538; and NHLBI T32HL83808. The contents of this manuscript do not represent the views of the U.S. Department of Veterans Affairs or the United States Government. Research reported in this publication was supported by the Strategic Health Innovation Partnerships (SHIP) Unit of the South African Medical Research Council (SA MRC) with funds received from the South African Department of Science and Technology as part of a bilateral research collaboration agreement with the Government of India; and through a SA MRC Collaborating Centre (ACT4TB/HIV). Additional support was also received through the Sub-Saharan African Network for TB/HIV Research Excellence (SANTHE), a DELTAS Africa Initiative [grant no. DEL-15-006]. The DELTAS Africa Initiative is an independent funding scheme of the African Academy of Sciences (AAS)’s Alliance for Accelerating Excellence in Science in Africa (AESA) and supported by the New Partnership for Africa’s Development Planning and Coordinating Agency (NEPAD Agency) with funding from the Wellcome Trust 107752/Z/15/Z and the UK government.

### Author Contributions

Conception and design – EWM, CLZ, MCG, DML; Acquisition of data – EWM, JGT, DIW, EK, SK, GS, SS, AW; Analysis and interpretation of data – all authors; Drafting of manuscript – EWM, DML; revisions and approval of manuscript – all authors

### Disclosure

The authors have no conflicts of interest to declare.

---

**Supplemental Table 1. Differential gene expression of lung derived MAIT cells and non-MAIT CD8^+^ T cells**

Genes that were enriched in MAIT samples from lung (compared to lung CD8^+^ non-MAIT) at thresholds of FDR-adjusted P value *≤* 0.001; and log2 fold change > 1 or <-1.

**Supplemental Figure 1.**
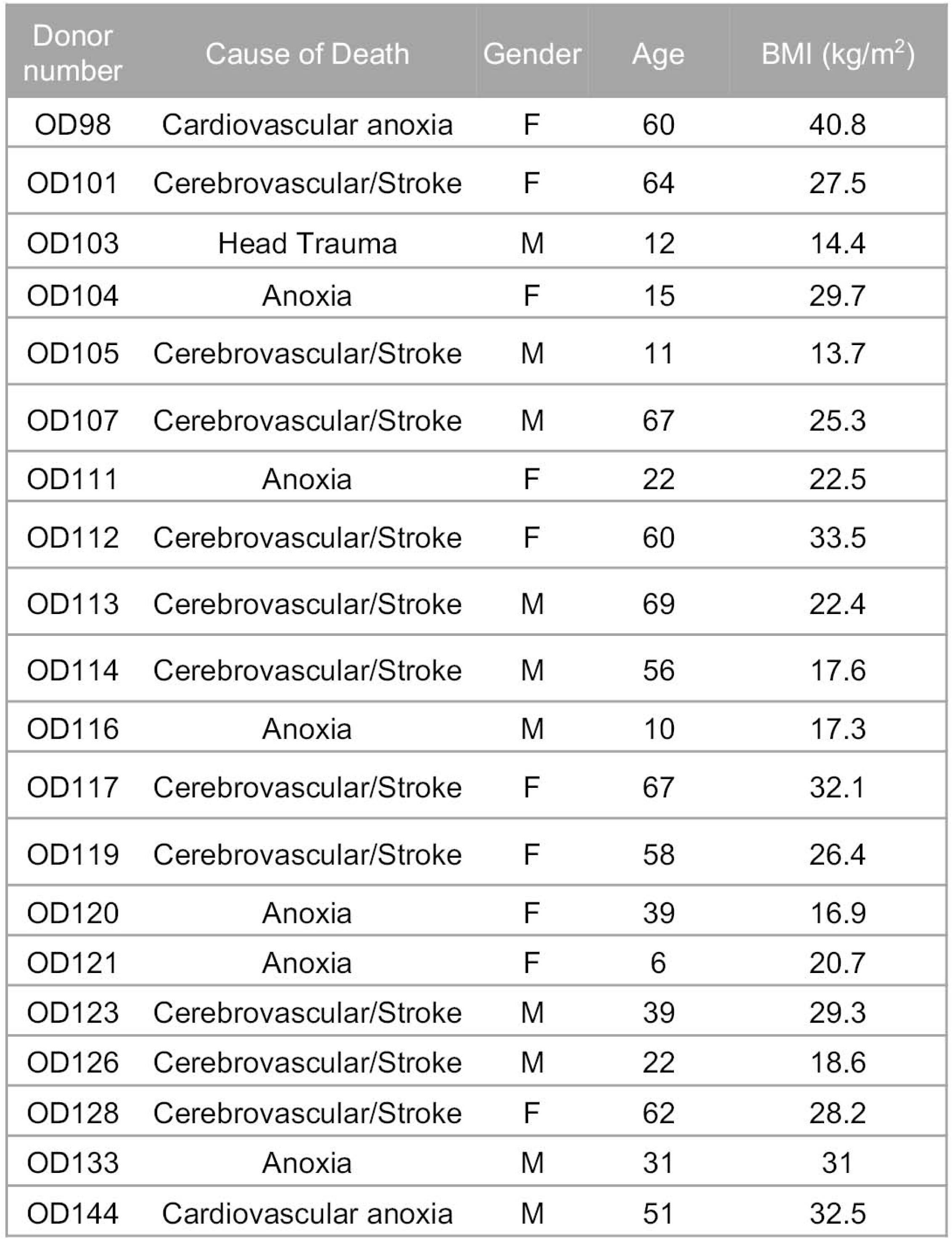
Pacific Northwest Transplant Bank donors provide lung tissue for single T cell analyses. Donor lung tissue was accepted for our study based on criteria detailed in the methods section. OD103, 104, 105, and 107 were used for RNA-seq analysis. The remaining lung donors were used for validation experiments.

**Supplemental Figure 2.**
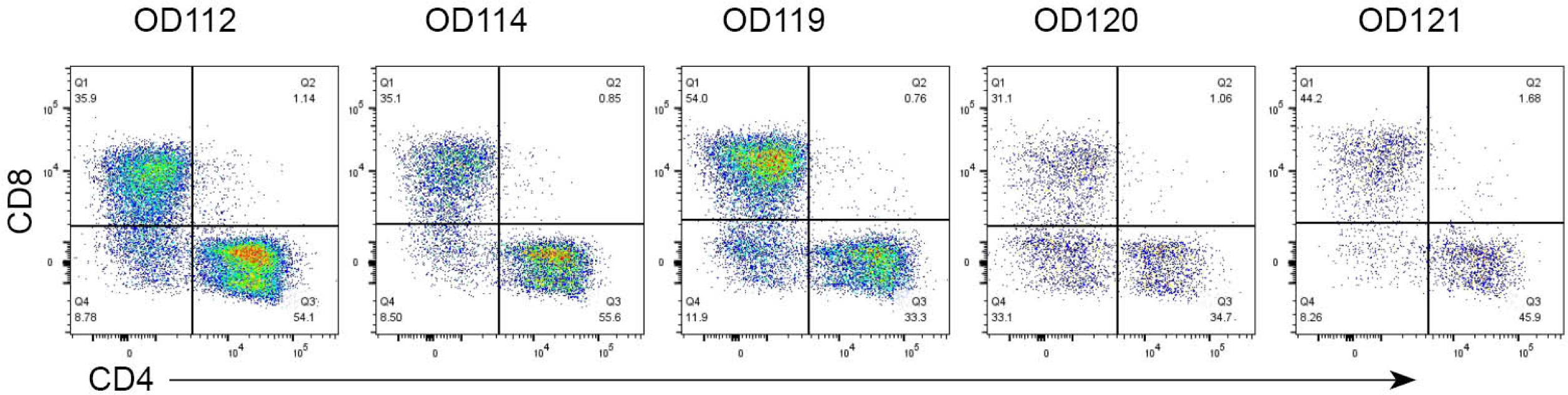
Lung MAIT cells express CD8 and CD4. Flow cytometry on lung samples to measure CD4 and CD8 co-receptor expression. Lung cells were rested overnight and expression on all CD3^+^ T cells are shown.

**Supplemental Figure 3.**
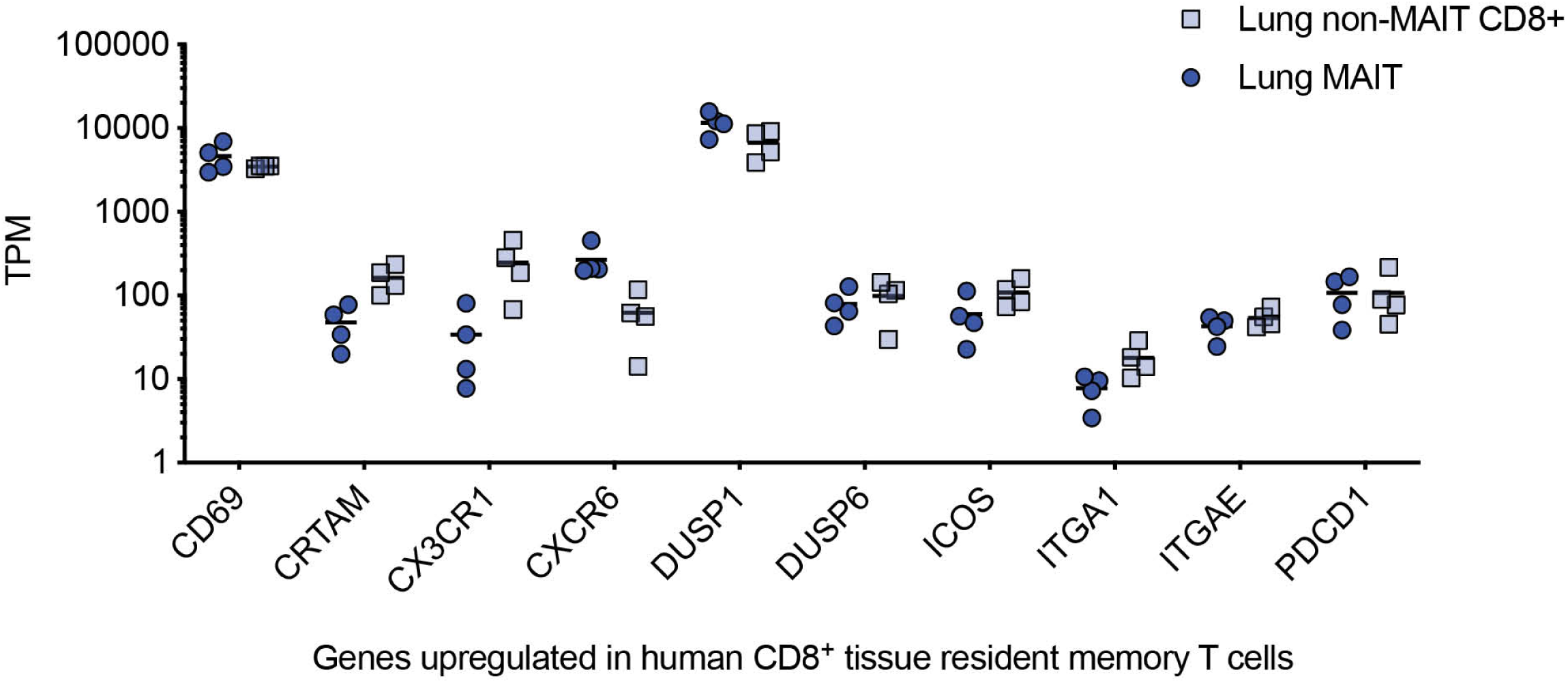
Lung MAIT cells express genes associated with human tissue-resident memory CD8^+^ T cells. Transcripts per million (TPM) reads from MR1/5-OP-RU^+^ CD8^+^ T cells derived from lung samples (Lung MAIT, blue filled circles), and MR1/5-OP-RU negative CD8^+^ T cells derived from lung samples (Lung non-MAIT CD8^+^, blue open squares). The mean of each group of samples is shown as a black bar. Genes listed on the x-axis are grouped by whether they have been observed as upregulated (A) or downregulated (B) in human CD8^+^ tissue-resident memory T cells compared to non-tissue resident memory T cells.

**Supplemental Figure 4.**
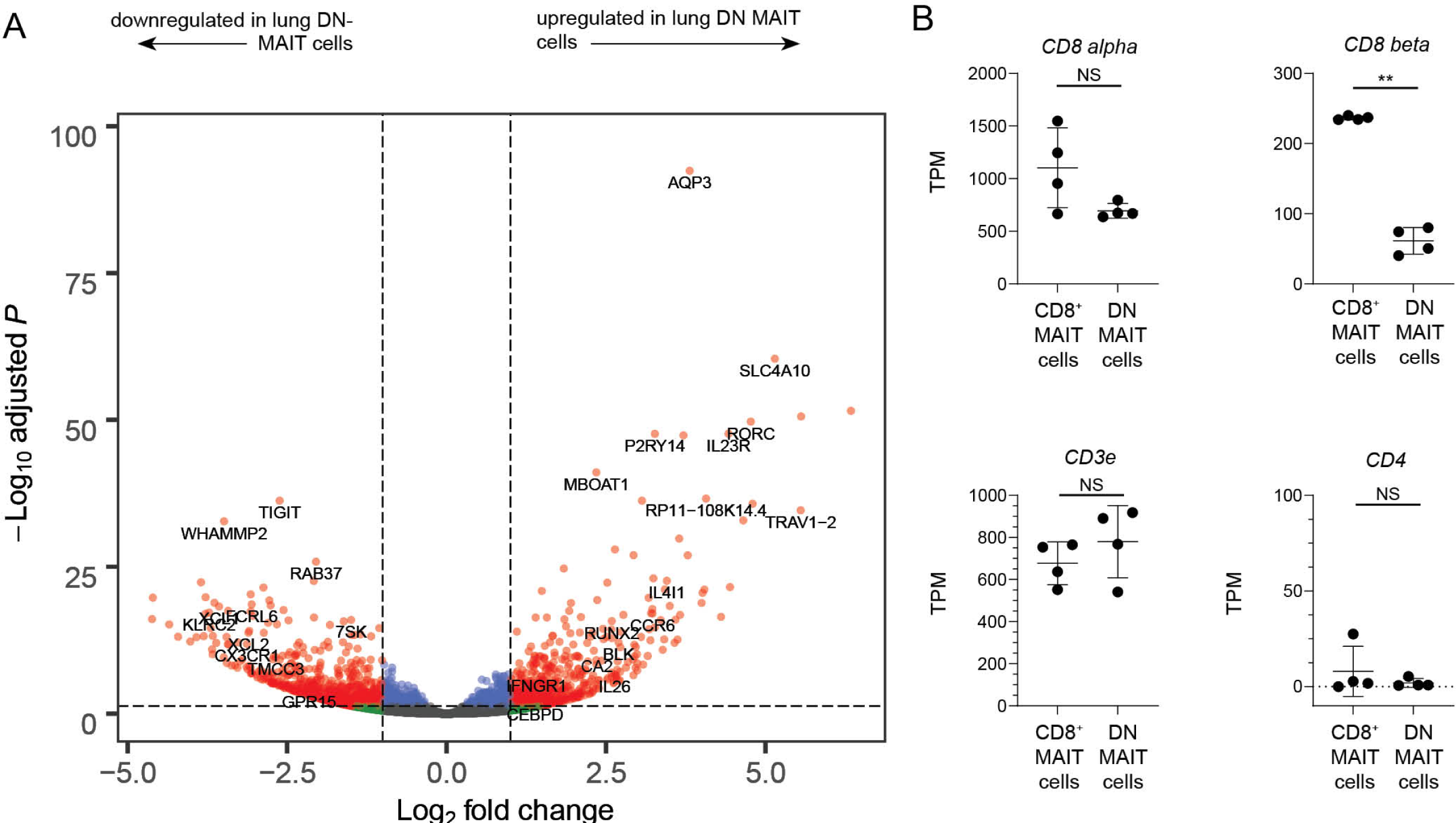
Differentially expressed genes of CD8^-^CD4^-^ MAIT cells in the lung. A. A volcano plot displays the differentially expressed genes between lung derived CD8-CD4-MAIT cells (on the right) and lung-derived CD8-CD4-lymphocytes (left) . Significantly differentially expressed genes (log2 fold change >1 or <-1, and FDR p-value *≤* 0.05) are represented as red. B. Box plots of transcript per million (TPM) reads of genes commonly associated with MAIT cells, from RNA-seq samples of CD8+ MAIT cells or non-MAIT CD8+ T cells derived from four lung samples. Box plots indicate the median and range of the TPM RNA-seq reads. FDR *P < 0.05, **P < 0.01, ****P < 0.0001.

**Supplemental Figure 5.**
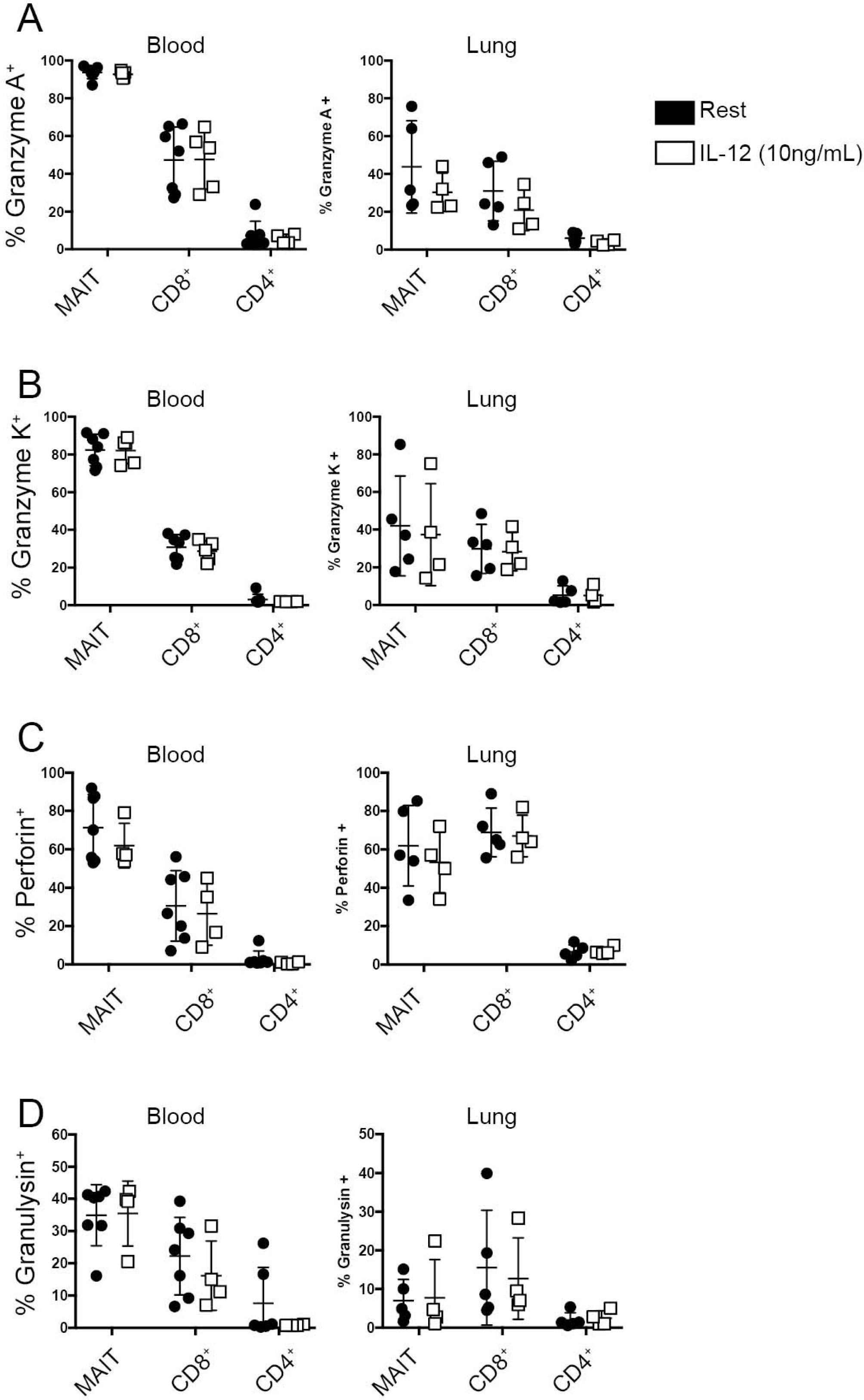
Poised expression of cytolytic proteins by subset of human T cells in the lung compared to the blood. Cells derived from PBMC or lung samples were either rested (black squares) overnight or in the presence of IL-12 cytokine (white squares, 10 ng/mL) and then stained intracellularly for (A) granzyme A, (B) granzyme K, (C) perforin, (D) granulysin. The frequency of cells staining positively for each protein is plotted from 5 PBMC donors (left) and 5 lung donors (right). Representative results of three experiments are shown, n = 5 biological replicates.

**Supplemental Figure 6.**
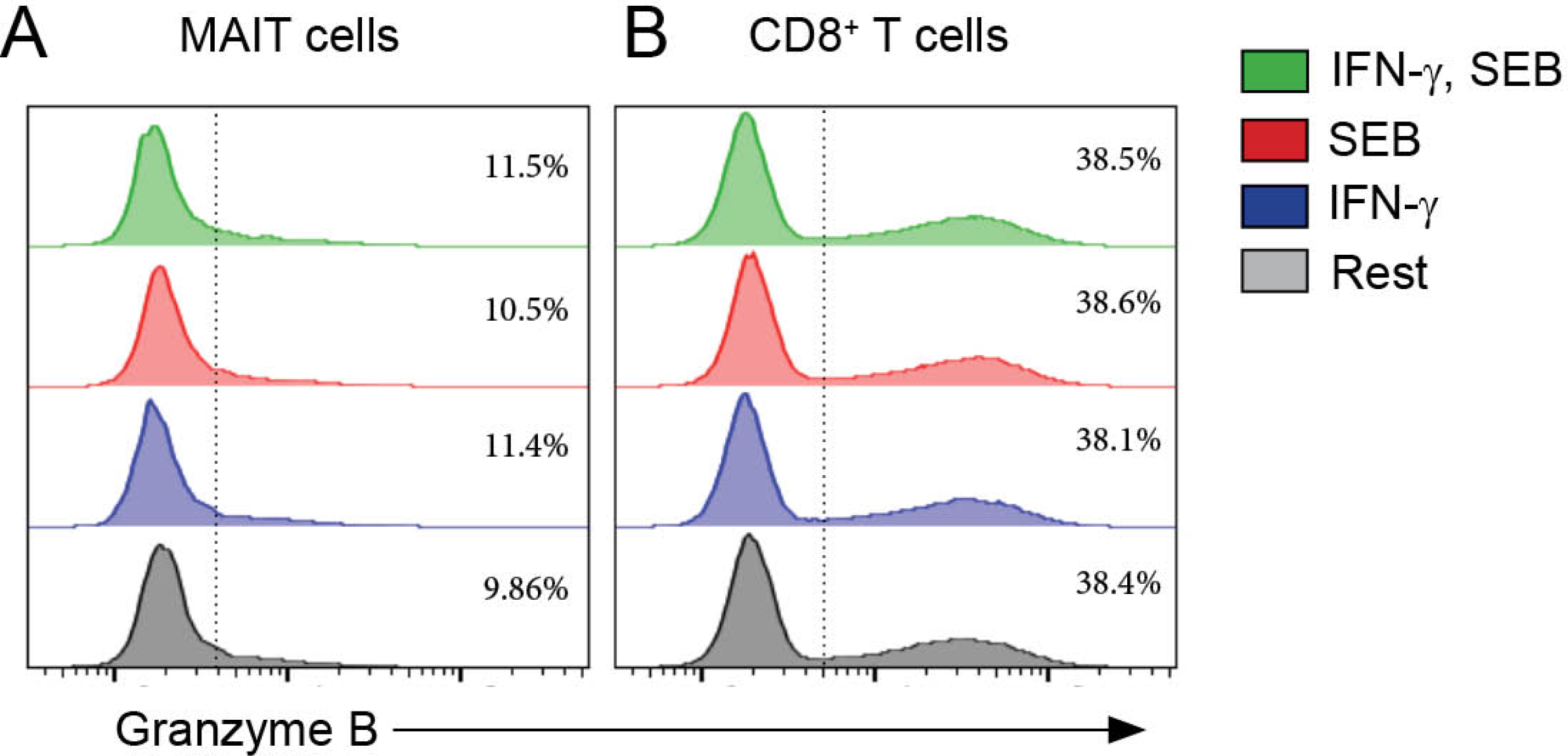
Intracellular levels of granzyme B in MAIT cells in the context of TCR stimulation or IFN-γ. Cells derived from PBMC were either rested (gray histograms) overnight or in the presence of IFN-*γ* (blue histograms, 10 ng/mL), SEB (red histograms), or IFN-*γ* and *Staphylococcus* enterotoxin B (SEB) (green histograms) and then stained intracellularly for granzyme B. The frequency of granzyme B^+^ MR1-tetramer^+^ MAIT cells (A) or non-MAIT CD8^+^ T cells (B) from one of five representative donors is plotted. Representative results of two experiments are shown.

