## Supplementary figures and images for "Human Lung-resident Mucosal-Associated Invariant T cells are Abundant, Express Antimicrobial Proteins, and are Cytokine Responsive"

### Supplemental Figure 1

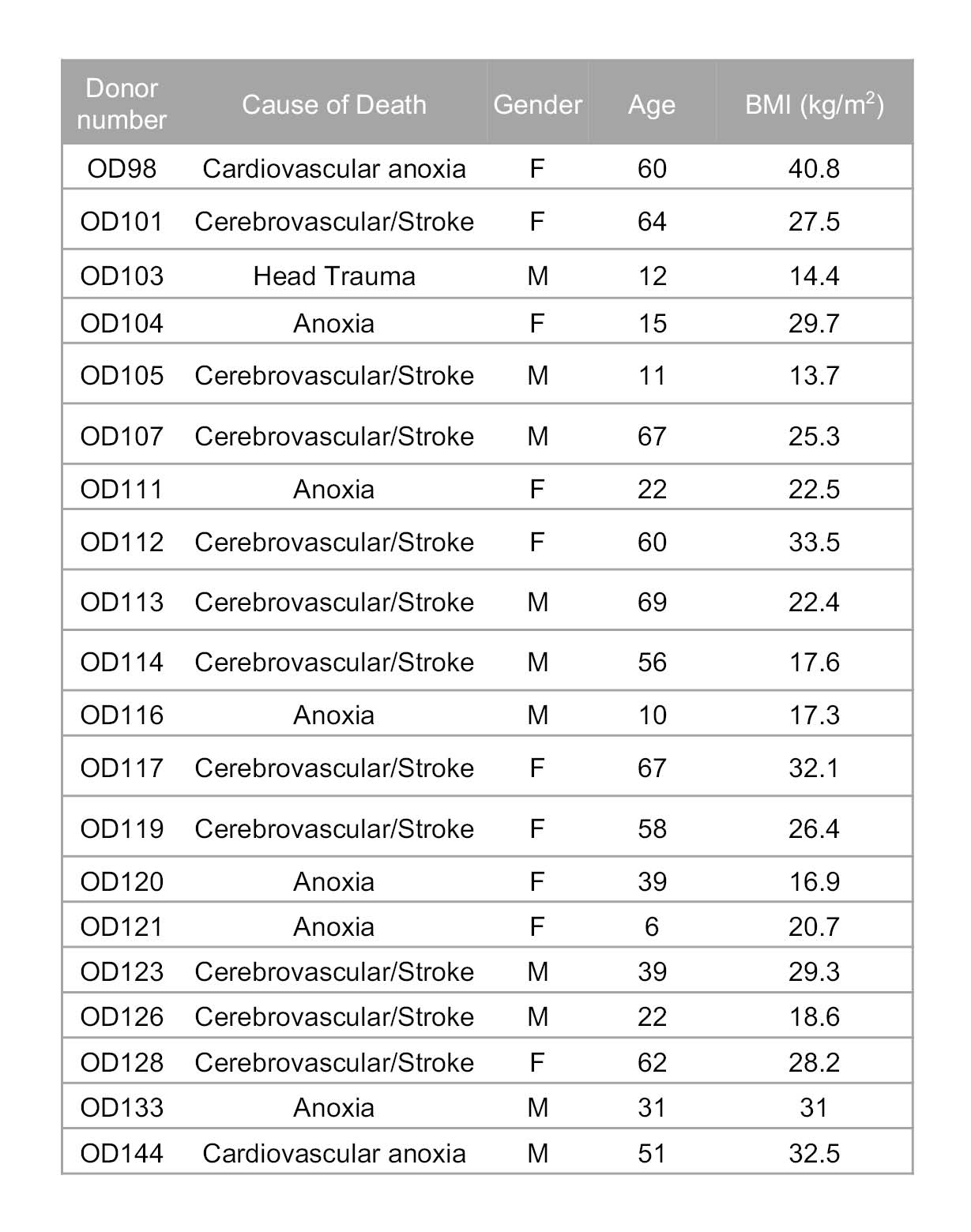

### Supplemental Figure 2

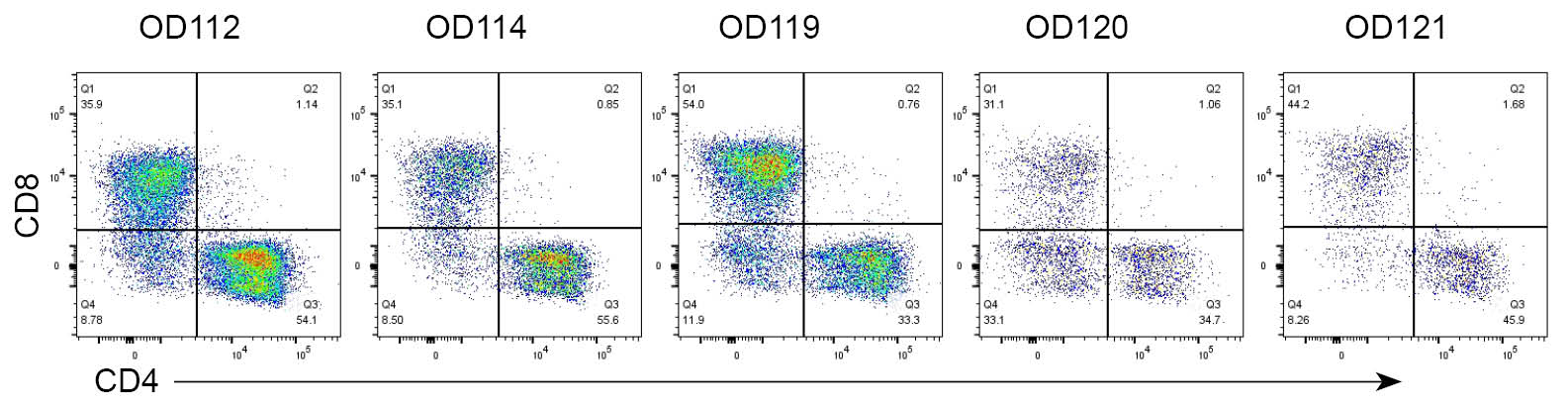

### Supplemental Figure 3

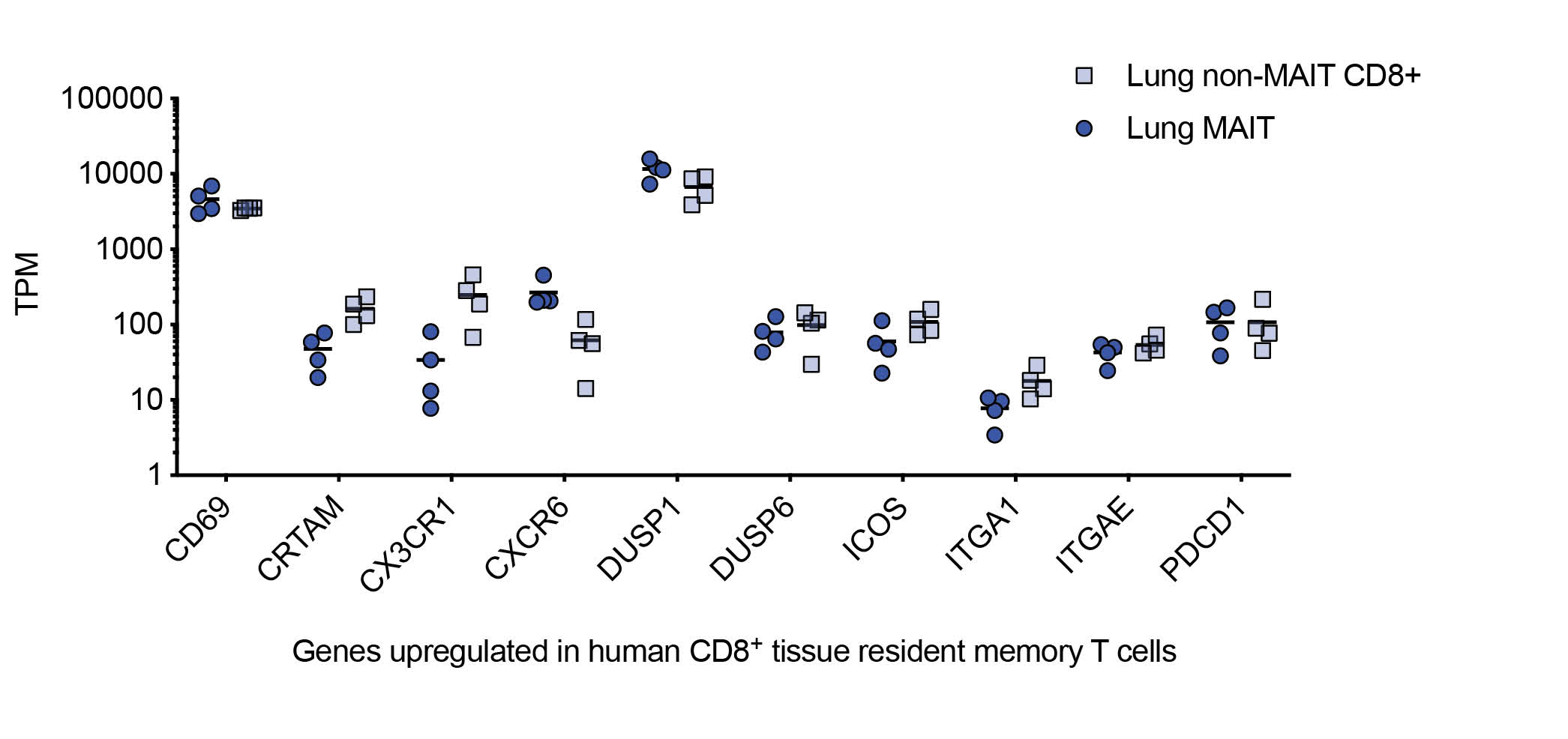

### Supplemental Figure 4

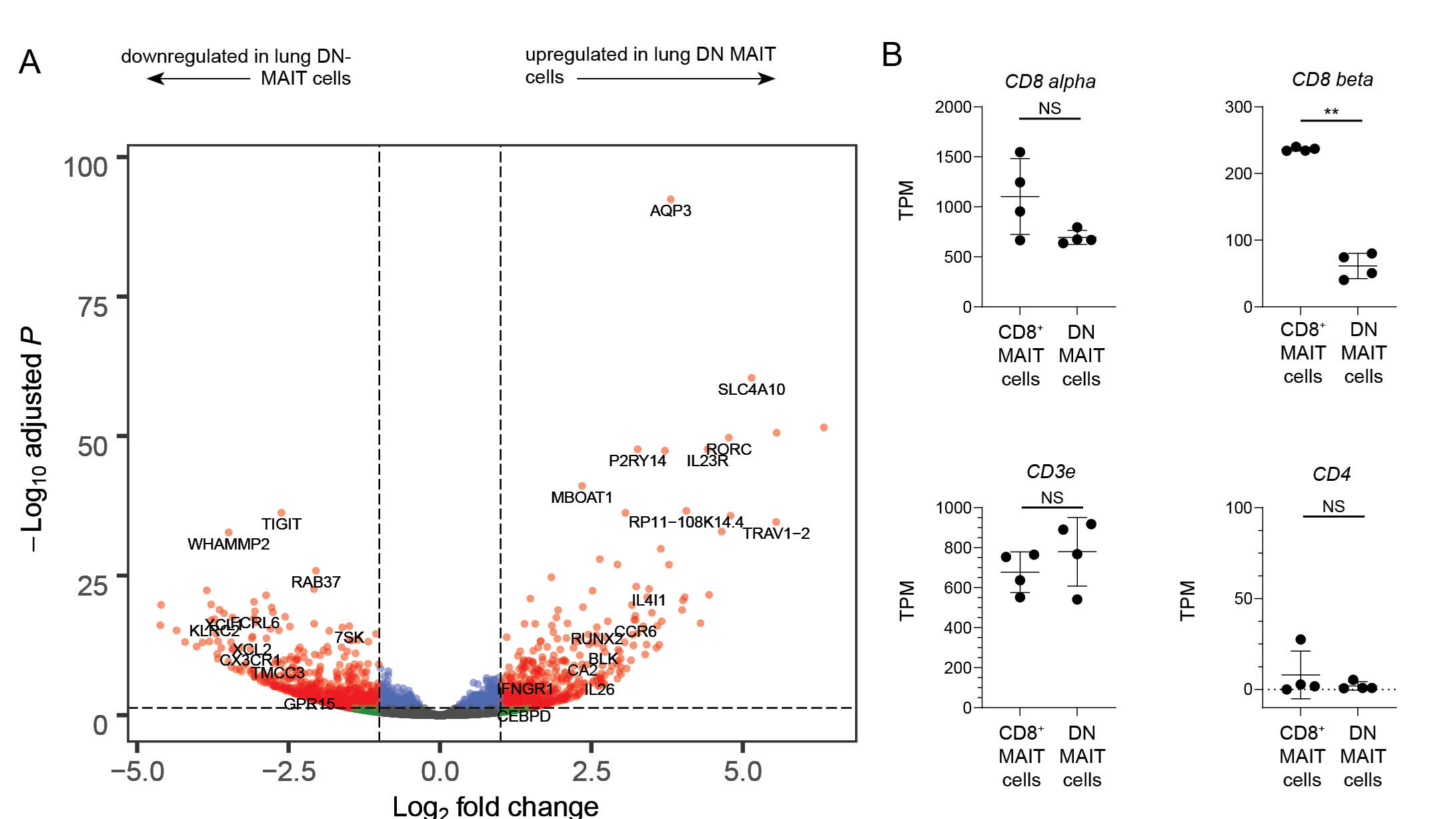

### Supplemental Figure 5

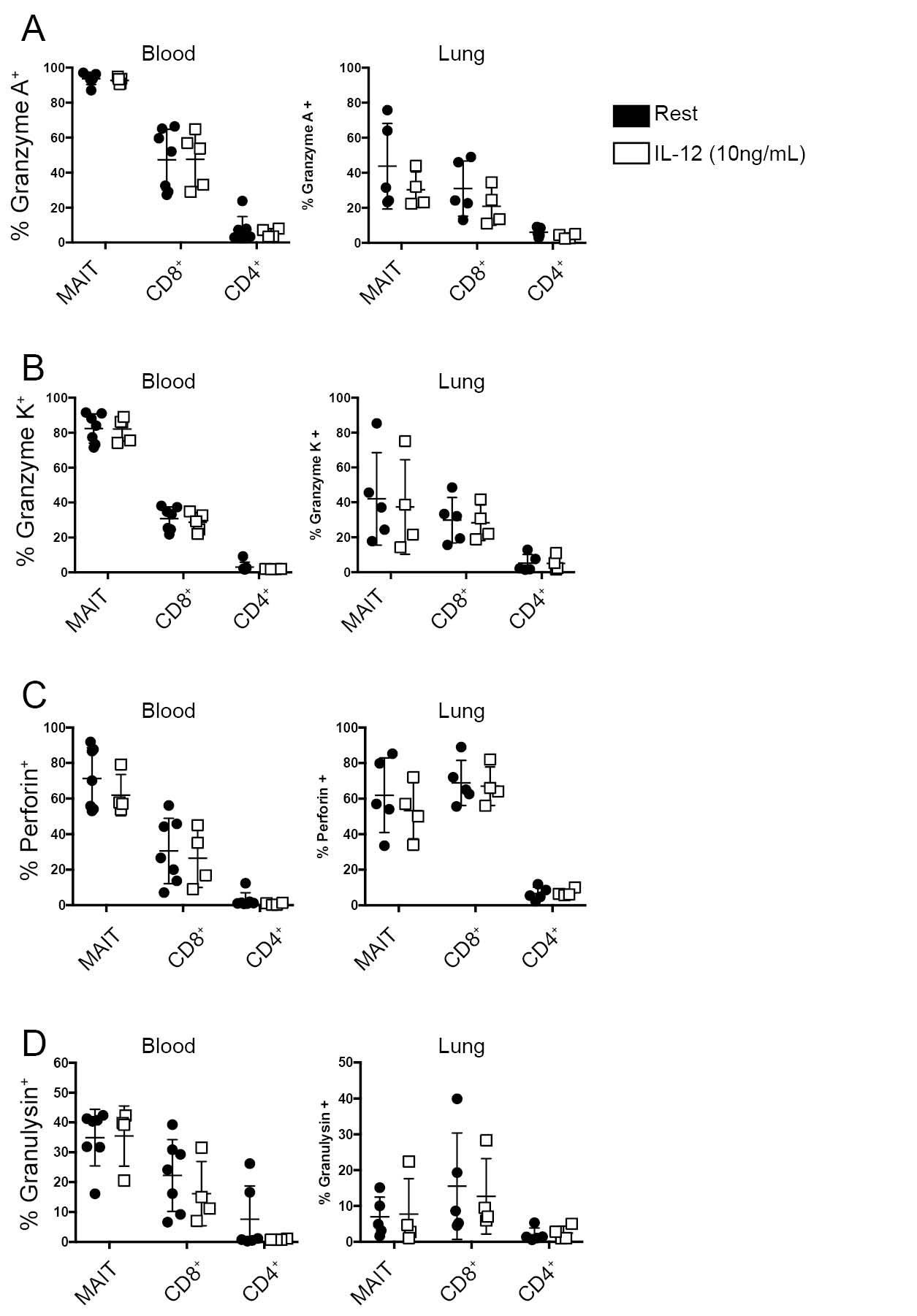

### Supplemental Figure 6

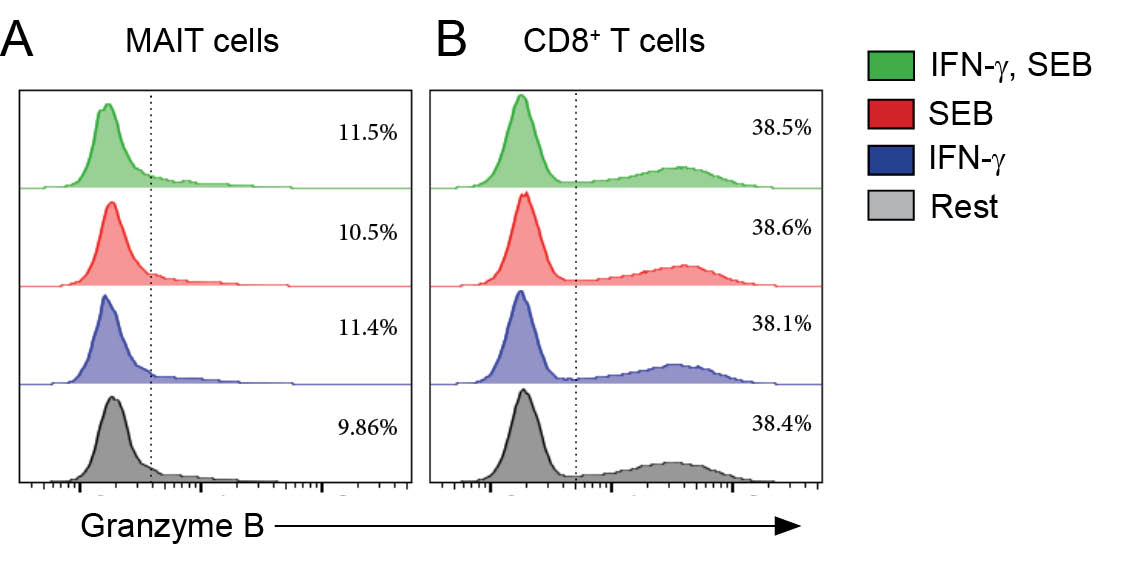
